# *In situ* structure and priming mechanism of the rhoptry secretion system in *Plasmodium* revealed by cryo-electron tomography

**DOI:** 10.1101/2022.01.11.475861

**Authors:** Matthew Martinez, William David Chen, Marta Mendonça Cova, Petra Molnár, Shrawan Kumar Mageswaran, Amandine Guérin, Audrey R. Odom John, Maryse Lebrun, Yi-Wei Chang

## Abstract

Apicomplexan parasites secrete the contents of rhoptries into host cells to permit their invasion and establishment of an infectious niche. The rhoptry secretory apparatus (RSA), which is critical for rhoptry secretion, was recently discovered in *Toxoplasma* and *Cryptosporidium*. It is positioned at the cell apex and associates with an enigmatic apical vesicle (AV), which docks one or two rhoptries at the site of exocytosis. The interplay among the rhoptries, the AV, and the parasite plasma membrane for secretion remains unclear. Moreover, it is unknown if a similar machinery exists in the deadly malaria parasite *Plasmodium falciparum*. In this study, we use *in situ* cryo-electron tomography to investigate the rhoptry secretion system in *P. falciparum* merozoites. We identify the presence of an RSA at the cell apex and a morphologically distinct AV docking the tips of the two rhoptries to the RSA. We also discover two new organizations: one in which the AV is absent with one of the two rhoptry tips docks directly to the RSA, and a second in which the two rhoptries fuse together and the common tip docks directly to the RSA. Interestingly, rhoptries among the three states show no significant difference in luminal volume and density, suggesting that the exocytosis of rhoptry contents has not yet occurred, and that these different organizations likely represent sequential states leading to secretion. Using subtomogram averaging, we reveal different conformations of the RSA structure corresponding to each state, including the opening of a gate-like density in the rhoptry-fused state. These conformational changes of the RSA uncover structural details of a priming process for major rhoptry secretion, which likely occur after initial interaction with a red blood cell. Our results highlight a previously unknown step in the process of rhoptry secretion and indicate a regulatory role for the conserved apical vesicle in host invasion by apicomplexan parasites.

## Introduction

*Plasmodium* parasites, the causative agents of malaria, were responsible for an estimated 627,000 deaths in 2020 with most deaths attributable to *Plasmodium falciparum*^1^. Despite significant progress in disease control over the last few decades, deaths increased by 12% from 2019^2^. *Plasmodium* spp. belong to a larger phylum of important disease-causing parasites called Apicomplexa, including *Toxoplasma* and *Cryptosporidium* that impact a large portion of the global population^3–5^. Invasive stages of these parasites, such as *Plasmodium* merozoites, contain a defining apical complex that harbors specialized structures and secretory organelles — micronemes and rhoptries — dedicated to parasite motility and host cell invasion^6, 7^. Micronemes first secrete their proteins onto the parasite surface, permitting parasite motility, host cell recognition, and downstream invasion events^6, 8^. Upon recognition of a host red blood cell, a pair of club-shaped rhoptries in the *Plasmodium* merozoite secrete their contents into the cytosol of the host, permitting the merozoite’s invasion and development within^7, 9^. The mechanism by which rhoptries secrete their contents, and how proteins from inside the rhoptries are relocated across the rhoptry membrane, parasite plasma membrane, and host plasma membrane remain largely unknown^10^.

Earlier electron tomography studies of merozoites undergoing red blood cell invasion demonstrated that the two rhoptries fuse together following host contact coupled with a reduction in volume, indicating secretion of contents into the host cell^11^. Similarly, traditional electron microscopy of merozoites invading red blood cells revealed that the merozoite’s two rhoptry necks are fused together^12^, or only one rhoptry (likely a fully fused rhoptry pair) is present^9, 13, 14^, with electron-translucent regions. Moreover, a pore has been seen at the interface between the fused rhoptry tips and the parasite plasma membrane (PPM), likely to provide a portal through which rhoptry proteins can relocate^13, 15^. The red blood cell surface protein glycophorin A (GlyA) has been demonstrated to distinctly trigger rhoptry secretion in isolated *P. falciparum* merozoites, however the mechanism of how this occurs remains unknown^16^. Recently, multiple non-discharge (Nd) proteins were discovered in *T. gondii* and *P. falciparum* that are implicated in the mechanism by which rhoptries secrete^17^. Calcium is likely involved in rhoptry fusion and secretion, as NdP1 and Fer2, two proteins containing calcium-binding C2 domains, are involved in rhoptry secretion and have been localized to the apical tip in *T. gondii*^17^. Moreover, fluorescence microscopy has revealed a calcium spike at the extreme apex of the merozoite just prior to penetration of the red blood cell^18, 19^. Despite these observations, however, the molecular mechanism by which this process occurs remains unclear.

Recently, cryo-electron tomography (cryo-ET) of *T. gondii* and *C. parvum* has revealed an elaborate molecular assembly, called the rhoptry secretory apparatus (RSA), that docks the rhoptries and an apical vesicle (AV) atop the rhoptries’ tips to the parasite apex^17, 20^. Knockdown of the Nd9 protein disrupted the RSA structure in *T. gondii* and abolished rhoptry secretion in both *T. gondii* and *P. falciparum*, linking the Nd-NdP proteins, the RSA, and rhoptry secretion together. However, whether *P. falciparum* harbors an RSA and/or AV at the rhoptry tips has yet to be determined. Furthermore, if the AV was genuinely part of this unique eukaryotic secretion system, then this would imply that there are two passages (one between the rhoptry and AV; another between the AV and PPM) that need to open for rhoptry secretion. The evidence for, purpose and regulation of the passages are unclear since there have been scarce hints as to what the role of the AV might be^10^.

Here we performed high-resolution cryo-ET imaging of isolated *P. falciparum* merozoites to investigate the ultrastructure of their rhoptry secretion system. We consistently observed an RSA in all the imaged merozoites and an AV in nearly two-thirds of them. We further resolved the RSA in detail through subtomogram averaging, a process that improves resolution by aligning and averaging multiple copies of the same structure in 3-dimensions (3-D). The elaborate structure of the RSA is similar to that of *T. gondii* and *C. parvum*^20^, all of which anchor into the PPM and form connections to the AV held below. Furthermore, we resolved two additional distinct states of the rhoptry secretion system that lack the AV: 1) with one of the two rhoptries directly docked at the RSA; and 2) with the two rhoptries fused together and the common tip docked at the RSA. Analyses of rhoptry volumes and rhoptry lumen densities revealed no significant changes across the three morphological states, suggesting that major rhoptry secretion has not occurred. We furthermore resolved different RSA conformations corresponding to each state through classified subtomogram averaging analyses, revealing multiple structural changes including the opening of a gate-like density in the RSA. Altogether, the different states likely reveal sequential events in rhoptry secretion, beginning with the opening of the passage between the AV and one of the two rhoptries, followed by fusion of the two rhoptries and gate opening in the RSA, suggesting a role for the AV in regulating rhoptry fusion to prime their secretion.

## Results

### Cryo-ET reveals an apical vesicle at the rhoptry tips in P. falciparum

Using cryo-ET, we imaged the apical region of isolated *P. falciparum* merozoites (Fig. 1a, b). We were able to resolve all the defining features of the apical complex, including two preconoidal rings, the apical polar ring, the inner membrane complex (IMC), rhoptries, micronemes, and subpellicular microtubules (Fig. 1b and Supplementary Fig. 1a-d). In addition, we observed a distinct AV atop the tips of the two rhoptries, whose presence has not been reported in *Plasmodium* (Fig. 1b, c) and is a distinct compartment from the two rhoptries that are docked just below, as evidenced by the significantly lower electron density within the AV versus the rhoptries (Fig. 1d, e and Supplementary Fig. 1e, f). As is the case with *T. gondii* and *C. parvum*, the AV in *P. falciparum* merozoites is docked at the PPM via the RSA (see details below). However, the AV of *P. falciparum* is smaller and its size and docking parameters are more variable compared to *T. gondii* and *C. parvum*^20^ (Fig. 1f-k and Supplementary Fig. 2). Of the 39 AVs analyzed, 34 (87%) had two rhoptries docked, four (10%) had just one rhoptry docked, and one (3%) had no rhoptry docked (Fig. 1l and Supplementary Fig. 1g-i). Of the AVs with two docked rhoptries, one rhoptry is always more aligned to the AV’s long (major) axis than the other (referred to as rhoptry 1 and rhoptry 2, respectively; Fig. 1g, i). In summary, our data reveal that the AV is a conserved feature across many apicomplexan parasites, docking the rhoptry tips to the RSA at the PPM.

**Figure 1.**
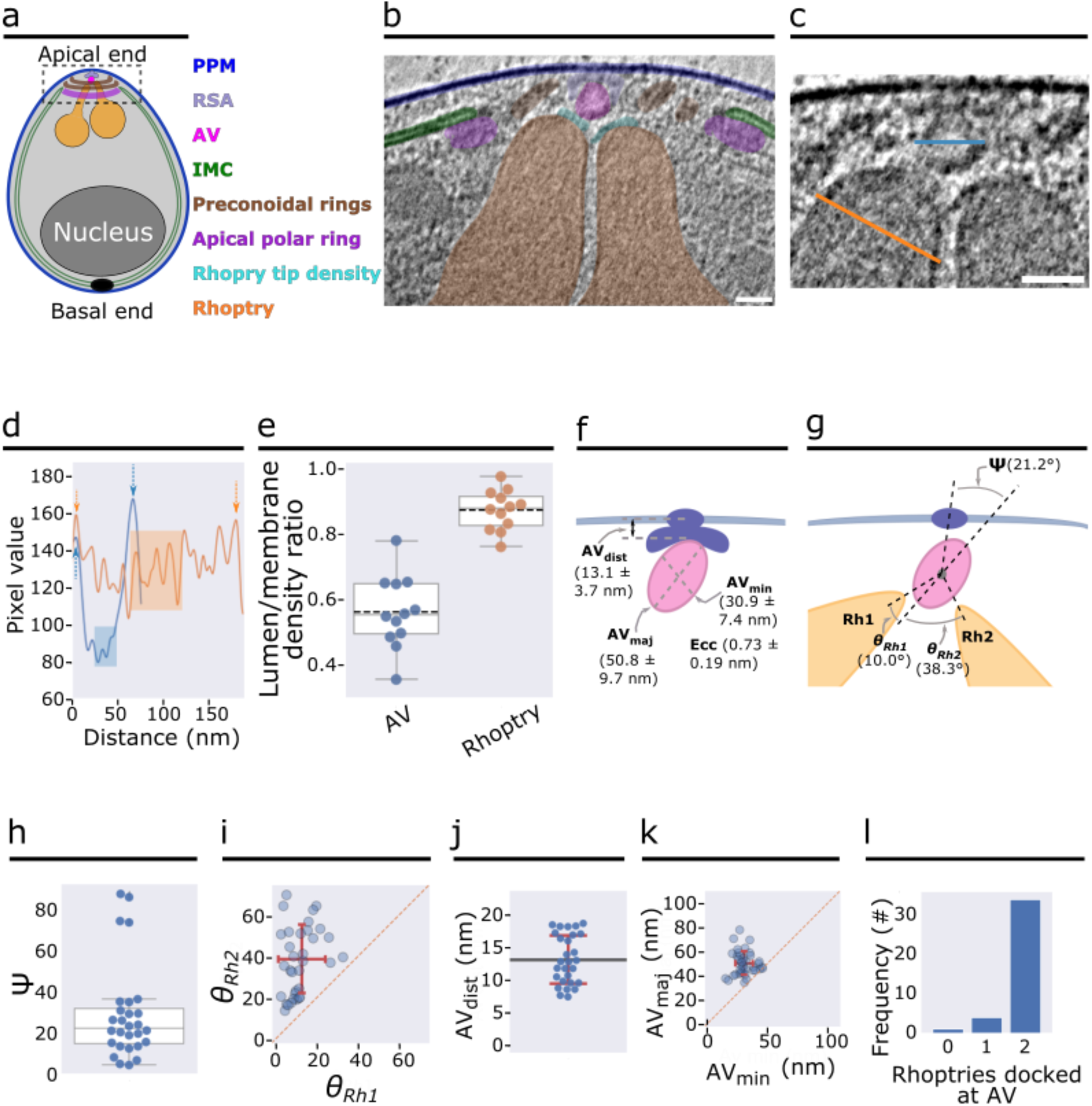
*P. falciparum* merozoites harbor an apical vesicle that docks the rhoptries to the parasite plasma membrane. **(a)** Simplified schematic of a *P. falciparum* merozoite. **(b)** 2-D slice from a tomogram of the merozoite apical complex displaying two rhoptries (orange), rhoptry tip density (cyan) two preconoidal rings (brown), an apical polar ring (purple), the parasite plasma membrane (PPM; dark blue), the inner membrane complex (IMC; green), the apical vesicle (AV; pink), and the rhoptry secretory apparatus (RSA; light blue). **(c)** Zoom-in view of the merozoite apex from panel b without color overlays. The orange and blue lines crossing the rhoptry and the AV, respectively, denote the locations of the pixel values shown in panel d. **(d)** Representative line profile plot of the inverted pixel values across the rhoptry (orange) and AV (blue). Arrows point to peaks corresponding to membranes. Shaded regions are the central 1/3 of pixels used to calculate the luminal density for the ratios shown in panel e. **(e)** Box plot showing the ratios of the luminal density to the corresponding membrane density (averaged) for each AV (blue) and docked rhoptry (orange). Mean value represented by the dotted black line. N = 12 AVs and 12 rhoptries. **(f)** Summary schematic of AV dimensions and docking distance to the PPM. **(g)** Summary schematic of the docking parameters among the rhoptries, AV, and RSA/PPM. **(h)** Plot of the angle (Ψ) at which the major axis of the AV is oriented with respect to the line connecting the PPM apex to the AV centroid, measured from several cells. **(i)** Plot of the angles at which rhoptry one (θ*_Rh1_*) and rhoptry two (θ*_Rh2_*) are docked at the AV, with respect to the major axis of the AV. **(j)** Measurement of the shortest distance between the PPM apex and the AV membrane, and **(k)** Plot of the major versus minor axes of the AV. **(l)** Histogram of the number of rhoptries docked at the AV in each cell. N = 39 AVs for panels h, j-l and N = 34 AVs and 68 rhoptries for panel i. Scale bars = 100 nm.

### P. falciparum merozoites display multiple rhoptry fusion morphologies

In addition to the discovery of the AV in *P. falciparum* (Fig. 1), we observed other rhoptry secretion system morphologies in a subset of merozoite tomograms analyzed (69/218; Fig. 2). Among them, 33 (15%) lacked the AV and instead had one of the two rhoptries docked directly to the RSA (Fig. 2b and Supplementary Fig. 3a-c). Rhoptries from merozoites displaying this morphology were generally more twisted and contorted as compared to rhoptry pairs from merozoites with an AV. Additionally, 36 (17%) merozoites display fusing rhoptries; again, these parasites lack an AV and instead the fused rhoptry tip is docked directly at the RSA. The fusion between the rhoptries ranged from fusion just at the rhoptry tips (Fig. 2c) to complete fusion to the base of the rhoptry bulbs (Fig. 2d and Supplementary Fig. 3d-g). Importantly, in all cases with fused rhoptries, no AV was observed. This suggests that the disappearance of the AV precedes the fusion of the rhoptry pair.

**Figure 2.**
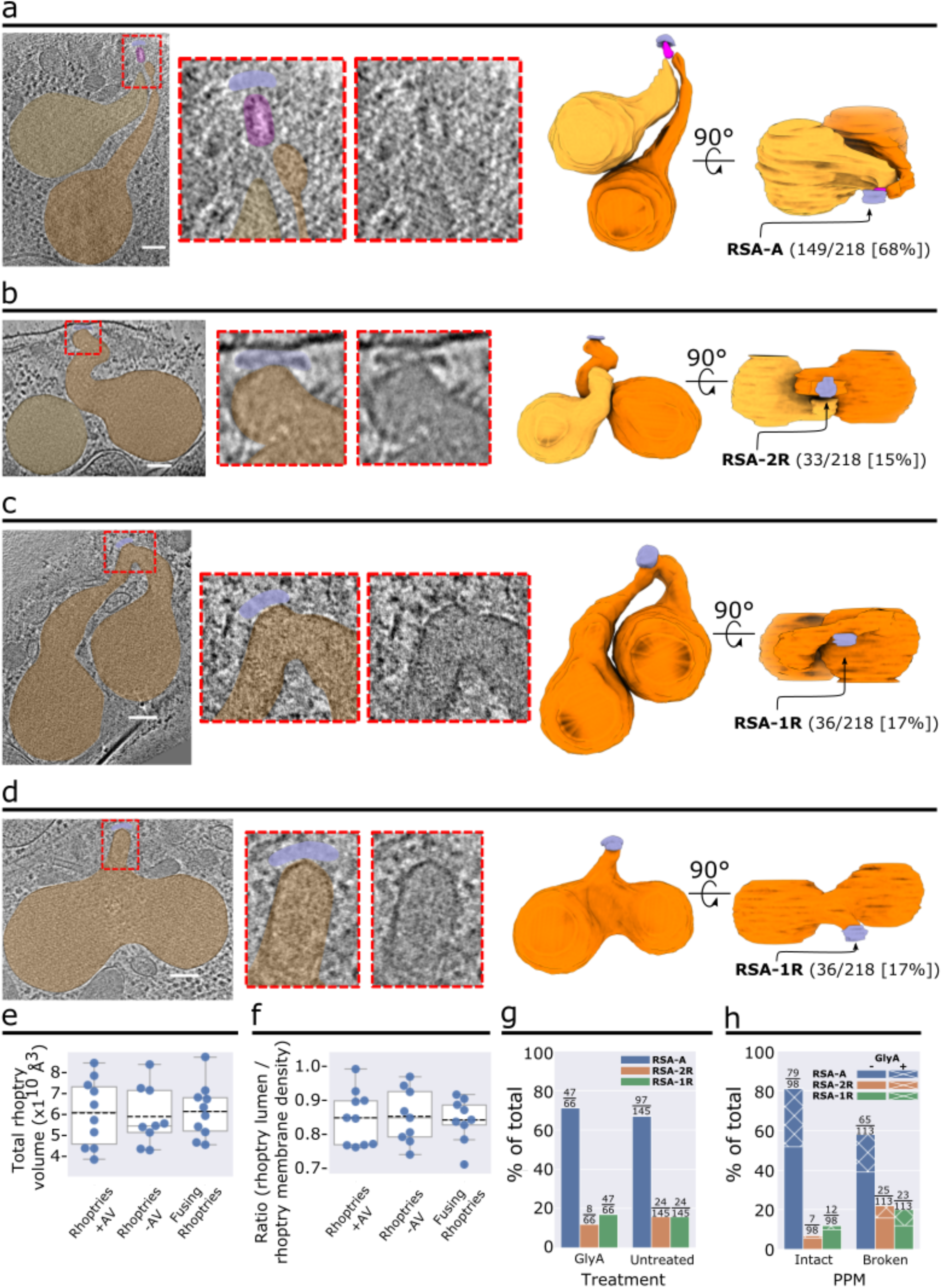
Various observed states of the rhoptry secretion system. **(a–d)** Representative examples of a merozoite apical end displaying two separate rhoptries docked at the AV (a), two separate rhoptries docked directly at the RSA (b), and rhoptries fused at the tips (c) and along the necks (d). Left: Tomogram slice of the apical end with color overlay on the RSA (light blue), AV (pink), and rhoptries (orange). Middle: Zoom-in of the apical tip of the rhoptries, with and without color overlay. Right: 3-D segmentations of the rhoptries, AV, and RSA with the proportion of analyzed merozoites exhibiting each morphology denoted. **(e)** Rhoptry volumetric analysis of 3-D rhoptry segmentations among the three rhoptry morphological states. N = 10 rhoptry pairs (left and right columns) and N = 9 rhoptry pairs (middle column). **(f)** Rhoptry luminal density analysis of 2-D slices of one rhoptry among the three rhoptry states from each tomogram analyzed in panel e. N = 10 rhoptries (left column) and 9 rhoptries (middle and right columns). **(g)** Plot of the proportion of *P. falciparum* merozoites, stratified by Glycophorin A (GlyA) treatment, and **(h)** Plot of the proportion of merozoites, stratified by PPM integrity, displaying each rhoptry morphology. Numbers above each bar represent the number of merozoites displaying each morphology over the total number of merozoites in each set (i.e., with or without GlyA treatment [g] or an intact or ruptured PPM [h]). N = 211 for total analyzed merozoites. Scale bars = 100 nm.

Rhoptry fusion has been previously observed during merozoite invasion of red blood cells when the rhoptries secrete their contents^11, 12^. We therefore sought to determine if the different rhoptry morphologies observed correspond to premature secretion even in the absence of host cells, although such an event is not expected during the merozoite’s normal invasion process. We identified 9-10 tomograms of each morphological class (separated rhoptry pair with an AV; separated rhoptry pair without an AV; or fused rhoptry pair without an AV) with entire rhoptries captured in the field of view. We then used EMAN2 neural networks^21, 22^ to segment the rhoptries in each tomogram and subsequently measured the volume of the rhoptry segmentations in UCSF Chimera^23^. The results show no significant difference in the average rhoptry volume among the different morphological classes (Fig. 2e). Similarly, density profiles of the rhoptry lumen also show no significant difference across the different classes (Fig. 2f). These observations suggest that there is no major secretion occurring upon rhoptry fusion in the absence of a host red blood cell. However, these data do not rule out the possibility of initial, small-scale secretions of specific rhoptry contents in preparation for a subsequent major secretion event.

In attempts to visualize rhoptry secretion and understand the triggers for these morphological changes, some of the merozoites were treated with GlyA prior to freezing on EM grids for cryo-ET imaging. We found that GlyA treatment did not affect the ratio of rhoptry morphologies observed (Fig. 2g), again supporting that these morphologies are not related to active secretion. Interestingly, we discovered that accidental rupture of the PPM (that likely happened during cryo-ET sample preparation) caused a significant decrease in the proportion of the morphology with an AV (from 81% to 58%; Fig. 2h). This suggests that the AV’s disappearance may be regulated by signals whose accessibility to the rhoptry secretion system can be influenced by the integrity of the PPM.

### Subtomogram averaging reveals an elaborate structure of the P. falciparum RSA

In all tomograms of merozoite apical complexes containing an AV, we found that the AV was always anchored to the PPM via the RSA (Figs. 1b, c, 2a, and Supplementary Fig. 1). A rosette of 8+1 intramembranous particles has been reported to be part of the *T. gondii* and *C. parvum* RSAs^20^ and linked to rhoptry secretion^17^. A similar rosette, albeit at lower resolution, has been previously reported by freeze-fracture electron microscopy at the apex of *Plasmodium* merozoites^17, 24^ and other apicomplexan parasites^25, 26^ as well. Resolving the overall RSA structure in *P. falciparum* merozoites will therefore be crucial to our understanding of the apicomplexan rhoptry secretion system and, in particular, species-specific adaptation in *Plasmodium*. We performed subtomogram averaging using all the RSAs associated with an AV (RSA-A) to boost signal over noise to analyze structural details of this large molecular assembly. The resulting average of the *P. falciparum* RSA revealed a density of much greater complexity than what was observed in individual tomograms (Fig. 3, Supplementary Fig. 4b, c, and Movie 1). As part of the RSA, we resolved an apical rosette of 8 membrane anchors (Anchor-I [A-I]) surrounding a central density (CD) at the PPM (Fig. 3b, c[i], and d[top]). Densities extend from each A-I towards the AV (Anchor-II [A-II] and Anchor-III [A-III]; Fig. 3b), showing a clear 8-fold arrangement in the intracellular space (Fig. 3c, d[side], f). The A-II connects to A-I in the PPM (Fig. 3b), and A-III connects to the AV (Fig. 3c[v] and Supplementary Fig. 5). Furthermore, in the center of the 8-fold A-II/III densities lies a central channel (CC) that sits atop the tip of the AV (Fig. 3b[iii], c[iii], and e). The CC connects to another set of 8 anchors (Anchor-IV [A-IV]; Fig. 3b[ii], c[i]) in the PPM via the radiating spokes (rS; Fig. 3c[ii], e). We also resolved a density at the interface of CC and the AV, termed the gate density (gD; Fig. 3b[iii], e[bottom], f[bottom]). All the structural details revealed by subtomogram averaging of the *P. falciparum* RSA docked with the AV are summarized in 3-D in Movie 2.

**Figure 3.**
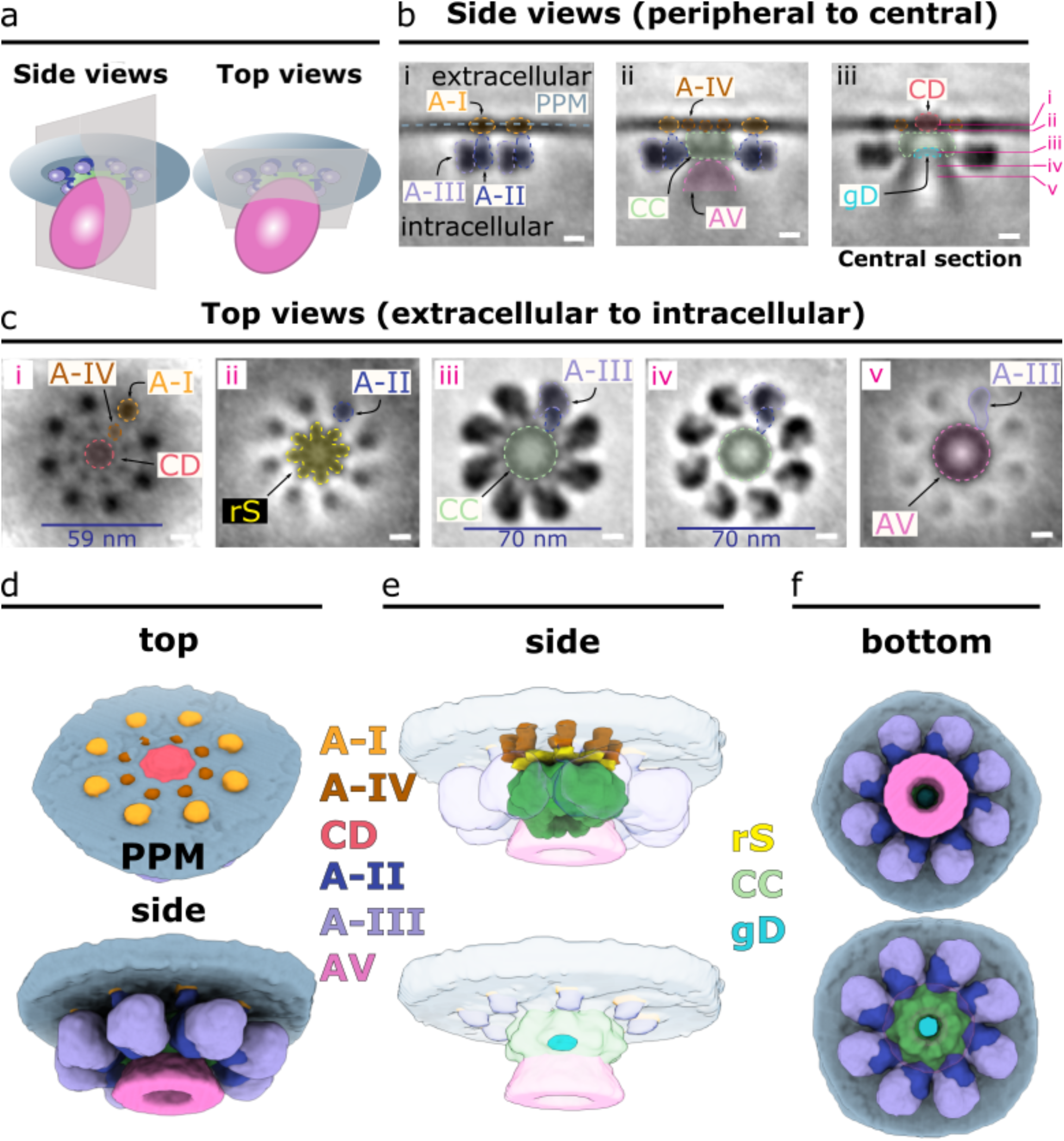
*In situ* structure of the multicomponent rhoptry secretory apparatus of *P. falciparum*. **(a)** Schematic of the RSA, AV, and PPM demonstrating the side view and top view orientations used to generate 2-D slices of the subtomogram average in panels b and c. **(b)** Side view slices of the RSA-A subtomogram average, from peripheral (left) to central (right) cross sections. The corresponding cross sections of the top view slices in panel c are denoted by the pink lines in b(iii). Colored outlines and overlays denote components of the RSA observed in these views: The Anchors I-IV (A-I, A-II, A-III, A-IV), the central channel (CC), the central density (CD), and the gate density (gD). **(c)** Top view slices of the RSA subtomogram average, from extracellular (left) to intracellular (right) cross sections. Colored outlines and overlays denote components of the RSA observed in these views: the radiating spokes density (rS) and other components as labeled in panel b. **(d)** 3-D segmentation of the RSA displaying outer most components from top and side views. **(e)** 3-D segmentation of the RSA displaying interior components from side views. **(f)** 3-D segmentation of the RSA from a bottom view with the AV shown (top) and not shown (bottom) to reveal the gD (bottom). Scale bars in panels b and c = 10 nm.

### The RSA displays a conserved ultrastructure among apicomplexans

We next compared the subtomogram average of the RSA and AV of *P. falciparum* with that of *T. gondii* and *C. parvum* previously resolved by our group^20^. Side views of the three RSA structures show a similar organization: the RSAs are anchored to the PPM intracellularly and holds the AV just below (Fig. 4a). Consistent with the greater variability in size and positioning of the *P. falciparum* AV compared to that of the two other organisms (Supplementary Fig. 2), only a small portion of the AV closer to the RSA was resolved in *P. falciparum* (Fig. 4a[i]), while more of the AV and its additional connections to the RSA were resolved in *T. gondii* and *C. parvum* (Fig. 4a[ii], 4a[iii]). Additionally, the membrane bound by the RSA displays significant curvature in both *T. gondii* and *C. parvum* while remaining flat in *P. falciparum*. Top views at the PPM display a conserved rosette of particles in all three parasites, however *P. falciparum* appears to contain another inner rosette of A-IV particles (Fig. 4b). The CD in both *P. falciparum* and *C. parvum* is positioned at the PPM (Fig. 4a[i], b[i], a[iii], and b[iii]), while that of *T. gondii* extends into the extracellular space (Fig. 4a[ii], b[ii]). Additionally, the outer rosette of A-I particles appears differently shaped between the three parasites: in *P. falciparum* they are circular (Fig. 4b[i]), whereas in *T. gondii* they appear elongated (Fig. 4b[ii]) and in *C. parvum* they appear teardrop shaped (Fig. 4b[iii]). The top views of the A-I, the CC and the A-II/III densities reveal a conserved 8-fold organization with some differences in the shape of the anchors (Fig. 4b, c). The shapes of each component can be better understood with a 3-D view of each RSA (Fig. 4d). From this view, we observe a similar arrangement of A-I densities at the PPM in all three parasites (Fig. 4d[top row]) with differences in the extent to which they protrude from the membrane, as well as differences in membrane curvature. From the side views (Fig. 4d[bottom row]), we can see that both *P. falciparum* and *T. gondii* have relatively globular A-II/III densities with limited or flexible (therefore largely averaged out in the subtomogram averages) contacts with the AV (Fig. 4d[i, bottom], d[[ii, bottom]). On the other hand, the *C. parvum* RSA makes extensive, rigid contacts with the AV that are resolved in the average and contribute to the full inclusion of the AV in the final average (Fig. 4d[iii, bottom]). Additionally, the *C. parvum* RSA contains a posterior central channel that sits within the AV (Fig. 4d[iii, bottom]) that is not seen in both *P. falciparum* and *T. gondii* AVs. Together, we show that the three apicomplexan rhoptry secretion systems contain an ultrastructurally conserved secretory apparatus with notable species-specific alterations in the docking of the AV at the PPM.

**Figure 4.**
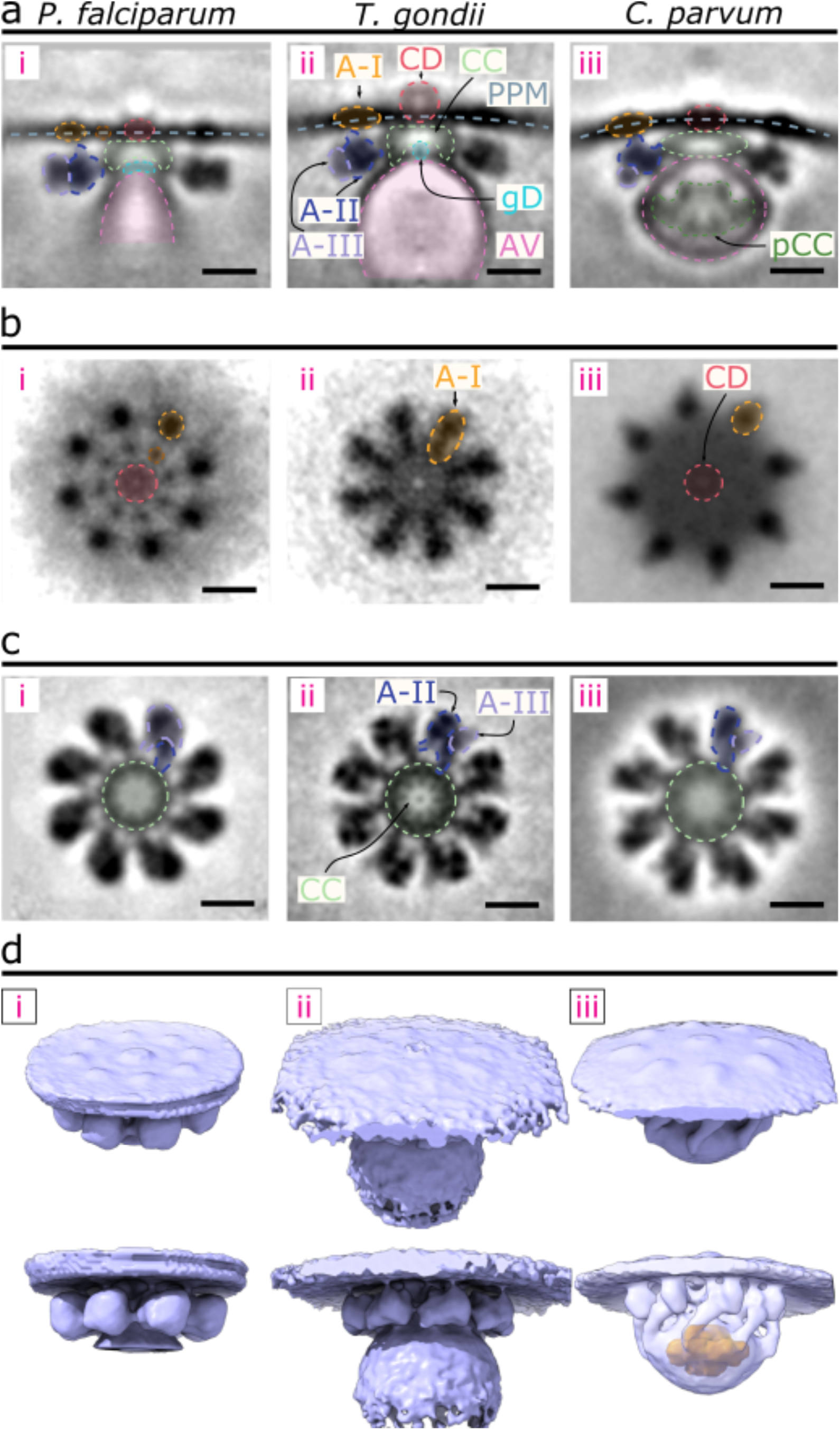
Comparison of the RSA structures of *P. falciparum*, *T. gondii*, and *C. parvum*. **(a)** Central slices of the side view of each RSA subtomogram average (left – *P. falciparum*; middle – *T. gondii*; and right – *C. parvum*) with color overlays of each corresponding component. **(b)** Top view of each RSA, comparing the densities observed on the PPM surface. **(c)** Top view of each RSA through a cross-sectional plane in the intracellular region, comparing CC, A-II, and A-III densities. **(d)** 3-D isosurface of each RSA, with views from outside the PPM (top row) and from the side (bottom row). The AV of *C. parvum* is made semi-transparent to reveal the posterior central channel density inside (orange). Scale bars = 20 nm.

### RSA structure changes with rhoptry fusion

To investigate if rhoptry fusion with the AV or fusion between the two rhoptries (Fig. 2) induces conformational changes in the RSA structure, we performed separate subtomogram averaging of the RSA particles classified under these different AV-rhoptry morphologies: those with an AV docked and two separate rhoptries (RSA-A; Fig. 2a and Movies 1, 2), those with no AV and two separate rhoptries (RSA-2R; Fig. 2b, Supplementary Fig. 4d-g, and Movie 3), and those with no AV and fusing rhoptries (RSA-1R; Fig. 2c, d, Supplementary Fig. 4h-k, and Movie 4). The central slice from the side view of each structure shows similar docking of the AV to the RSA (RSA-A) or of the rhoptry tip to the RSA (RSA-2R and RSA-1R), but with different membrane curvatures at the interface (Fig. 5a). Multiple RSA components appear different among the three structures. First, from the top view at the level of the PPM, there is a significant reduction of the CD and disappearance of the A-IV densities in the RSA-1R structure compared to the other two (Fig. 5b[iii]). Additionally, the diameter of the rosette of A-I particles decreases by ∼3.5 nm between the RSA-A and RSA-2R structures, and by another ∼1 nm between the RSA-2R and RSA-1R structures (Fig. 5b). At a slice through the CC, we observe significantly less electron densities within the channel of the RSA-1R structure compared with the other two (Fig. 5d). To ensure this finding is not due to contrast differences among the averages, we plotted the normalized pixel intensity across the CC from each structure and show a clear decrease in relative pixel intensities within the RSA-1R CC (Fig. 5d, e [blue line]). Remarkably, from the side views of the structures, we identified the main cause for such a density decrease: loss of the gD at the interface between the CC and the fused rhoptry tip (Fig. 5f). Similarly, consistent with the top views, the CD was largely reduced in the PPM of the RSA-1R structure from the side view (Fig. 5f). Finally, the side views of the A-II/III densities revealed a ∼3.5 nm shift of them towards the PPM in the RSA-1R structure compared with the RSA-A structure (Fig. 5g). This upward shift of the A-II/III densities is also reflected in the top view slice through the rS density (Fig. 5c): we observe only A-II densities around the rS in the RSA-A and RSA-2R structures, whereas both A-II and A-III densities are seen in the RSA-1R structure. Moreover, we observe an additional connection formed between the A-III and the PPM (termed Anchor-V [A-V]; Fig. 5g) while the A-II density connecting to A-I appears largely reduced (Fig. 5a, g). Altogether, the results revealed intricate conformational changes of the RSA related to the sequential AV and rhoptry fusion activities. The detailed structural differences are illustrated in 3-D in Movie 5.

**Figure 5.**
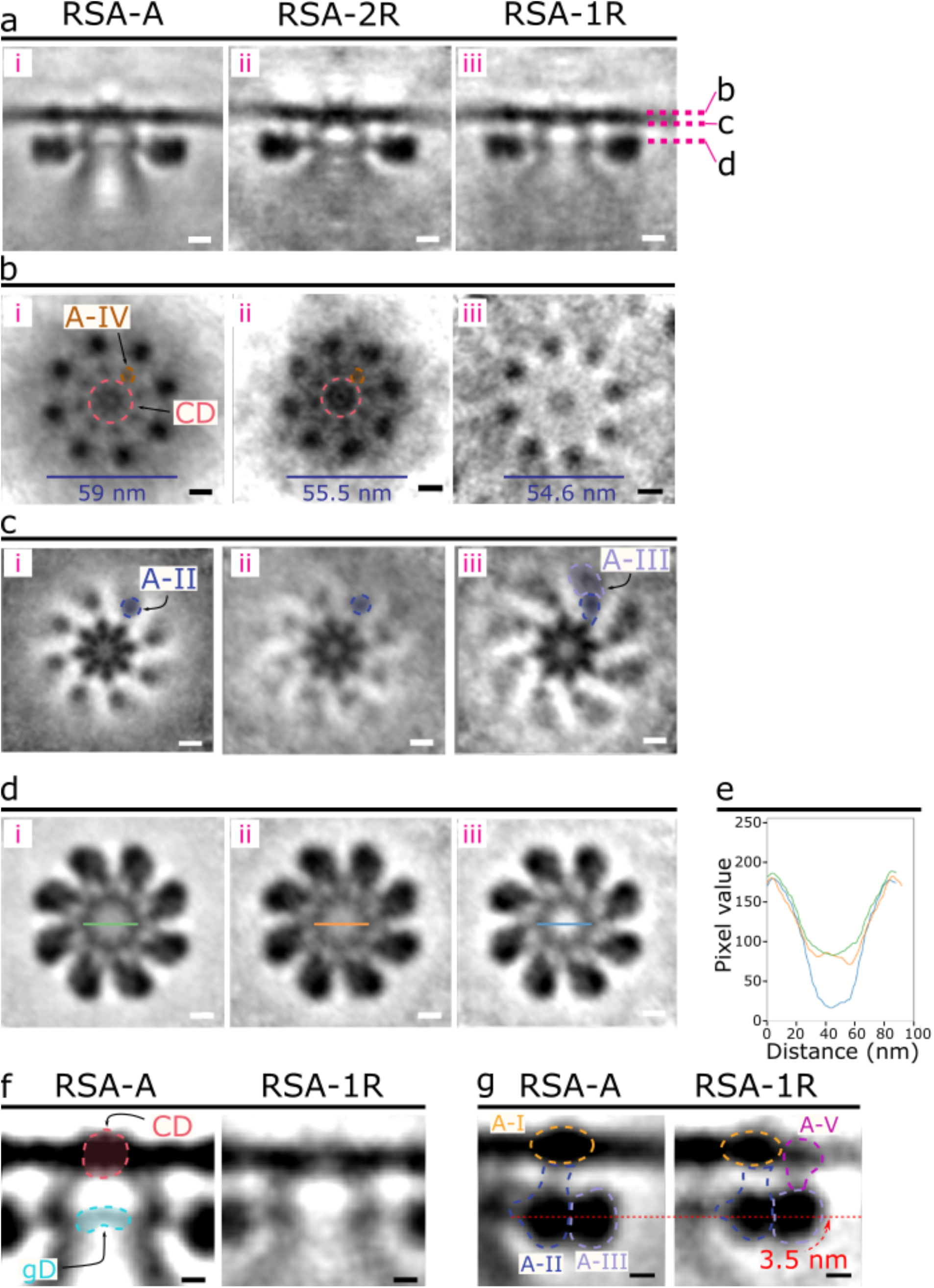
Different *P. falciparum* RSA conformations were resolved corresponding to different morphological states of the rhoptry secretion system. (a) Central slices of the side view of each RSA conformation (left – RSA-A; middle – RSA-2R; right – RSA-1R). Cross sections in panels b-d are denoted by the pink dashed lines. (b) Top view of each RSA conformation from outside the PPM. The CD and A-IV densities are observed in the RSA-A (i) and RSA-2R (ii) conformations but reduced in the RSA-1R conformation (iii). The diameter of the rosette formed by the A-I densities on the PPM surface is noted by the blue lines at the bottom. (c) Top view of each RSA conformation just below the PPM. A-II and A-III densities have color overlays to highlight their change in the RSA-1R conformation. (d) Top view of each RSA through an intracellular cross section. The green, orange, and blue lines across the CC of the RSA-A, RSA-2R, and RSA-1R, respectively, denote the location of the pixel values plotted in panel e. (e) Plot of normalized, inverted pixel values across the central channel of each RSA (RSA-A, green; RSA-2R, orange; RSA-1R, blue). A large dip is seen among the pixels of the RSA-1R CC. (f) Zoom in on panel a (i and iii) highlighting changes of the gD and CD between RSA-A and RSA-1R. (g) Zoom in on panel a (i and iii) highlighting changes in A-II/III between RSA-A and RSA-1R, including a diminished A-II connection to the A-I density and 3.5 nm upward shift of the A-II/III densities (curved red arrow), as well as the presence of a new Anchor-V (A-V) in the RSA-1R conformation. Scale bars = 10 nm in panels a–d and 5 nm in panels f, g.

## Discussion

### The AV is a conserved feature of the apicomplexan rhoptry secretion system

Previous cryo-ET imaging of *Plasmodium* revealed important insights into parasite motility and ultrastructure^11, 27^, however the information was limited due to thick samples and the inherent low contrast of biological samples. Cryo-ET is rapidly evolving and recent advancements in contrast enhancing transmission electron microscopy imaging technologies have allowed for the visualization of parasitic structures at molecular resolution in a native, frozen-hydrated state with cellular contexts^20, 28, 29^. For example, the use of contrast-enhancing phase plates and improved detectors have permitted novel insights into the conoid and rhoptry secretion system of invasive-stage *T. gondii* and *C. parvum* parasites^17, 20, 29^, while focused ion beam milling has allowed cryo-ET imaging of *P. falciparum*-infected red blood cells^30^. Here, we apply these technologies to image the apical complex of isolated *P. falciparum* merozoites and successfully resolve multiple defining features, including the rhoptry secretion system and multiple cytoskeletal components (Fig. 1, Supplementary Fig. 1a-d). Similar to previous reports^8, 11, 13, 31, 32^, we observe two rhoptries with necks that extend through the preconoidal rings (Fig. 1b). Unlike previous studies that have reported on the AV in *T. gondii*, *C. parvum*, and *Sarcocystis tenella*^20, 26^, no study has reported on this vesicle in *P. falciparum*. We report for the first time that the *P. falciparum* rhoptry tips indeed dock at a distinct AV in most merozoites (Fig. 1), similar to that of *T. gondii* and *C. parvum*^20^. The AV exhibits a small, ellipsoid shape with an electron-translucent interior. The rhoptry tips dock at the AV like in *T. gondii*, rather than being already fused to the AV membrane like that of *C. parvum* (Supplementary Fig. 2a). Whereas *C. parvum* sporozoites contains one rhoptry^33^, *P. falciparum* merozoites have two and *T. gondii* tachyzoites have 8-12^34, 35^ and thus docking at the AV, rather than being pre-fused, may represent a mode of regulation that permits secretion of multiple rhoptries at once. Furthermore, the presence of the posterior central channel within the *C. parvum* AV (Fig. 4a[iii], d[iii]) and the pre-fused rhoptry-AV membranes reflects the possibility that this organism exhibits an alternate mode by which it regulates fusion between the rhoptry tip and the AV. Alternatively, this may represent an intermediate stage with the same mode of secretion regulation that has already been partly activated in *C. parvum*. With this discovery of the AV in *P. falciparum*, we can now observe its presence in earlier cryo-ET images of merozoites^11, 30^. The AV in *P. falciparum* merozoites has likely been overlooked due to its small size and flexibility (Fig. 1 f-h, j, k and Supplementary Fig. 5) compared to that of other apicomplexan parasites^17, 20^. Recent comparative analyses between the apicomplexan rhoptry secretion system and the extrusome system in the related, free-living Ciliata phylum^36^ suggests that the AV is an adaptation in Apicomplexa for parasitism, likely involved in the invasion of a host cell^10, 17^. By resolving the apical vesicle in *P. falciparum* merozoites, we provide further evidence that this enigmatic vesicle is a conserved feature among apicomplexan parasites that may play a role in host cell invasion.

### Conserved RSA features suggest a similar activity and regulation of rhoptry secretion across Apicomplexa

Our recent report discovered the elaborate structures of the RSA in *T. gondii* and *C. parvum*, which exhibits complex interactions with the AV and the PPM to ultimately guide the rhoptries to the parasite apex via the AV (Fig. 4)^20^. Here, the extensive connections observed between the *P. falciparum* RSA and PPM and between the RSA and the AV (Fig. 3) again support the function of the RSA as one that docks the AV to the PPM; the AV in turn guides the rhoptries to the proper site for exocytosis. Detailed differences among the RSA structures from these organisms (e.g., the *P. falciparum* RSA contains two concentric membrane rosettes [A-I and A-IV densities], while *T. gondii* and *C. parvum* contain only one [Fig. 4b]) may reflect adaptations in the molecular mechanisms for their specific parasitic activities. Further cryo-ET imaging of these parasites in the context of host cell invasion will permit the species-specific dissection of RSA-mediated rhoptry secretion, thus, to understand the roles of each common or unique structural component of the RSAs. Identification of the molecular components that comprise the RSA is also necessary to further elucidate the mechanisms of secretion. The recent discovery of Nd proteins, which are necessary for rhoptry secretion in *T. gondii* and *P. falciparum* and may contribute to RSA function^17^, provide a hopeful avenue to begin probing RSA-mediated secretion by cryo-ET. For example, knockdown of the Nd9 protein abolishes rhoptry secretion and cryo-ET revealed this was due to altered RSA structure in *T. gondii*^20^. Further cryo-ET analysis of related mutants and/or conditions to map molecules onto the RSA structures will permit a more comprehensive understanding of this unique molecular machine.

### The AV may regulate rhoptry fusion to prime for secretion

Earlier electron micrographs of *Plasmodium* merozoites in the act of red blood cell invasion revealed the formation of a pore between the rhoptry membrane and the PPM^13, 37, 38^. Additionally, fusion between the two rhoptries during rhoptry secretion has been observed^11–13^. It has therefore been hypothesized that the rhoptries fuse together and with the PPM during secretion^39^, allowing the discharge of two rhoptries simultaneously. Our observation that the AV is only present at the tips of non-fused rhoptry pairs but always absent when the rhoptries were seen fused (Fig. 2a-d and Supplementary Fig. 3) suggests that the AV and its disappearing may participate in the regulation or initiation of rhoptry fusion in *P. falciparum*. A recent cryo-ET study of developing merozoites within schizonts showed images of the apical complex in which the AV was present at the rhoptry tips (Supplementary Fig. 1j)^30^, but was overlooked and instead marked as part of the rhoptry (Supplementary Fig. 1k, l). This indicates that AV biogenesis occurs during parasite development within the red blood cell, and that the rhoptry morphologies observed in merozoites lacking an AV are likely subsequent states to the AV-containing state. Moreover, in the morphology that lacks the AV while the two rhoptries are separated, the two rhoptry necks often twist around each other or display bulges that extend towards one another (Fig. 2b, Supplementary Fig. 3a-c). This may be indicative of the initiation of fusion between the two rhoptries, hinting at a role for the AV’s disappearance in triggering this activity. Indeed, similar observations of possible rhoptry fusion can be observed in earlier electron micrographs of *T. gondii* during host invasion, where the secreting rhoptries display branched morphologies that may be representative of two rhoptries fused together along the necks^34, 35, 40^. Although it has not been proven if the rhoptries are indeed fused in these data, the fact that no AV could be observed in any of the invading *T. gondii* parasites again supports a role of the AV’s disappearance in rhoptry function during host invasion.

Interestingly, volumetric and density analyses revealed that no major secretion occurs across all the rhoptry morphologies captured in our cryo-ET data (Fig. 2a-f), thus uncoupling rhoptry fusion from the major secretion event. This suggests that the fusion between the two rhoptries could be an independent event upstream of major content secretion and likely has its own functional purpose. The different RSA conformations revealed by subtomogram averaging for each rhoptry morphology further showed molecular rearrangements at the rhoptry exocytosis site correlating with the AV and rhoptry fusion activities. Importantly, the gD is absent, the CD is largely reduced, and the CC interior is less electron dense in the rhoptry-fused state (RSA-1R; Fig. 5e, f), seemingly to clear up the most direct path for rhoptry contents to leave the merozoite. Together, these data reveal a previously unknown step prior to secretion of rhoptry contents that may function as a priming mechanism to prepare the merozoite for major rhoptry secretion upon host contact. Despite no significant reduction in rhoptry volume or luminal density during this priming step (Fig. 2e, f), the possibility of small-scale secretions of specific components required for red blood cell invasion cannot be excluded. Indeed, previous studies have demonstrated that proteins important for invasion, such as Rh5, translocate from the rhoptries to the merozoite surface prior to full rhoptry secretion^31, 41^. Furthermore, the trigger for this priming step remains unknown as GlyA-treated parasites (GlyA was previously shown to be a trigger for rhoptry secretion^16^) displayed similar proportions of merozoites with and without the AV (Fig. 2g). Given the intimate link between calcium signaling and rhoptry secretion^16, 18, 19, 42^, the priming activity of rhoptries captured in our samples may be due to changes in intracellular calcium concentrations as the proportion of merozoites with primed rhoptries (the morphologies that lack the AV) increases with PPM rupture that likely causes entry of extracellular calcium into the cell (Fig. 2h). In accordance with this, various proteins required for the secretion of rhoptries are linked to calcium, including Rasp2^43^, Fer2, and NdP2^17^. Further analysis is necessary to confirm whether these fusion events are linked to calcium.

In summary, cryo-ET imaging of isolated *P. falciparum* merozoites has revealed a priming mechanism for the secretion of rhoptries (Fig. 6), in which the enigmatic AV may serve as a trigger to initiate rhoptry fusion. We have further resolved multiple RSA structures that correspond to the disappearance of the AV and fusion of the rhoptries, providing a structural framework to understand the mechanism at the molecular level. Through comparative analysis of the *P. falciparum* RSA to those of *T. gondii* and *C. parvum*, we strengthen the support for the AV’s role in host invasion by apicomplexan parasites in addition to evidence for conservation of this novel secretion system across Apicomplexa. Discovery of proteins required for rhoptry secretion that localize to the AV and RSA will permit the mapping of components onto this molecular machine *in situ* for fine dissection of the secretion mechanism that will provide novel targets for the treatment of this deadly parasite.

**Figure 6.**
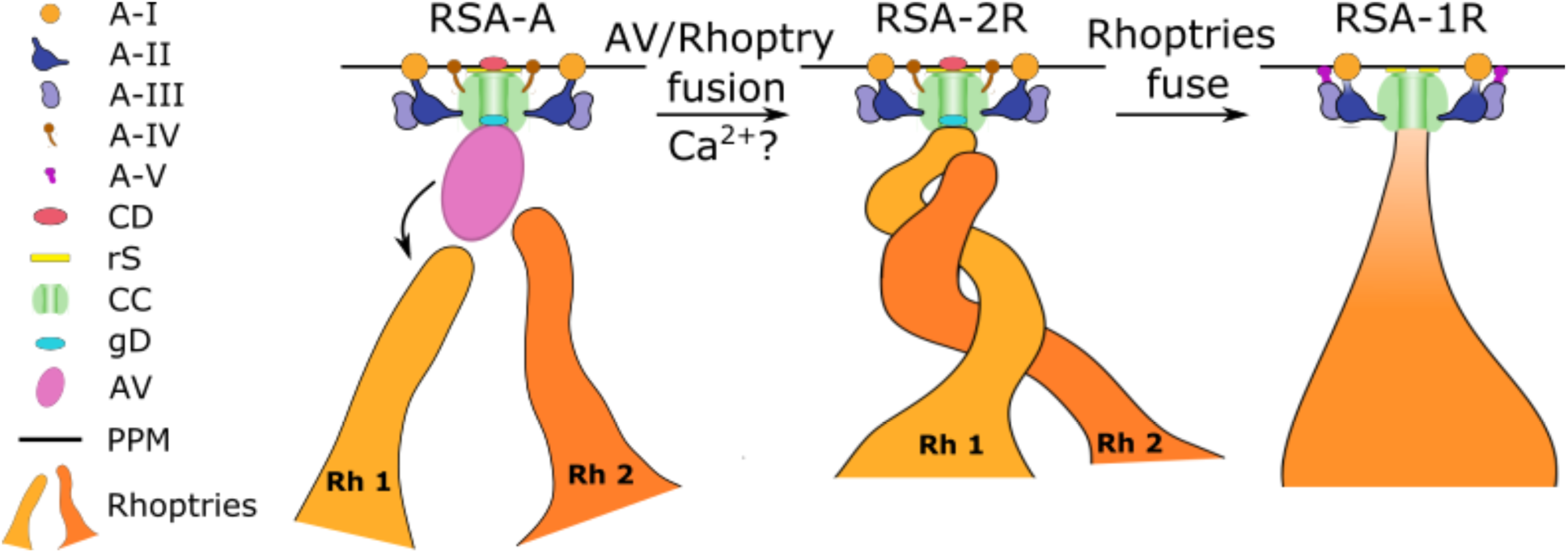
Working model of the priming of rhoptry secretion in *P. falciparum* merozoites. Prior to invasion signals, the AV docks to the RSA and guides the two rhoptries to their proper exocytosis site in preparation for secretion. Upon exposure to proper (unknown) signals, the AV fuses with one rhoptry (labeled Rh 1 for rhoptry 1), a process that may be mediated by calcium, creating the first passage opening for rhoptry secretion (i.e., between the rhoptry and AV). One rhoptry then docks directly at the RSA and prepares to fuse with the second rhoptry (labeled Rh 2 for rhoptry 2). During this process, the diameter of the A-I rosette in the PPM decreases and the membrane adopts a curvature around the CD. The rhoptries then fuse together partially (along the necks) or fully, resulting in a loss of the gD and a second passage opening, between the rhoptry and the RSA. This is accompanied by a further decrease in A-I rosette diameter, loss of the A-IV densities, formation of the A-V densities connecting the A-III densities to the PPM, and a large reduction of the CD density.

## Methods

### Culture of *P. falciparum* merozoites

*P. falciparum* 3D7 parasites were maintained at 37°C in a gas mixture of 5% O_2_, 5% CO_2_, and 90% N_2_ in a 2% suspension of human erythrocytes (donated from the Children’s Hospital of Philadelphia) in complete medium (RPMI-1640 completed with 27 mM NaHCO_3_, 11 mM glucose, 5 mM HEPES, 0.01 mM thymidine, 1 mM sodium pyruvate, 0.37 mM hypoxanthine, 10 μg/mL gentamicin (all aforementioned culture media components were purchased from Sigma Aldrich), and 5 g/L Albumax (Thermo Fisher Scientific)). Parasite cultures were synchronized over 3-4 cycles prior to merozoite isolation. Ring-stage parasite cultures were incubated with 5% (w/v) sorbitol (Sigma Aldrich) for 5 min at 37°C. Schizont-stage cultures were centrifuged in a 70% (v/v) isotonic Percoll (Sigma Aldrich) solution and the late-stage parasites were collected after treatment.

### Preparation of *P. falciparum* merozoites and cryo-ET grid preparation

For merozoite preparation, cultures were scaled up to 60 mL at 5% hematocrit. Late-stage schizonts were purified from highly synchronous cultures by magnetic activated cell sorting (MACS). Isolated schizonts were treated with 10 μM E64 for 6-8 hours, followed by passage through 1.2 μm syringe filters (preincubated in 1% BSA + PBS for 20 minutes) to mechanically isolate merozoites^44^. Isolated merozoites were spun down at 2200 *g* for 15 minutes and resuspended in 20 μL complete media. 10 nm colloidal gold fiducials (Ted Pella, Redding, USA) were added to the suspension (for alignment during tomogram reconstruction from tilt series). 4 μL of suspended merozoites were applied onto Quantifoil 200 mesh copper R2/2 holey carbon EM grids, excess liquid blotted away, and plunge frozen into a liquid ethane/propane mixture (pre-cooled with liquid nitrogen) using an EM GP2 automatic plunger (Leica Microsystems, Wetzlar, Germany)^45^. The blotting chamber was set to 95% relative humidity at 24°C and blotting was done from the sample side of the grid using Whatman filter paper #1. Plunge-frozen grids were subsequently loaded into autogrid c-clip rings (Thermo Fisher). The autogrid box containing frozen grids were stored in liquid nitrogen and maintained at ≤–170°C throughout storage, transfer, and cryo-ET imaging.

### Drug treatment of *P. falciparum* merozoites

For treatment with glycophorin A, mechanically isolated merozoites (prior to plunge freezing) were resuspended in buffer mimicking extracellular ionic conditions (140 mM NaCl, 5 mM KCl, 1 mM CaCl_2_) containing 1 mg/mL glycophorin A (Sigma Aldrich) and incubated at 37°C for 5 – 15 minutes^16^.

### Cryo-electron tomography (cryo-ET)

Cryo-ET was performed on a Thermo Fisher Krios G3i 300 keV field emission cryo-transmission electron microscope. Dose-fractionated imaging was performed using the SerialEM software^46^ on a K3 direct electron detector^47^ (Gatan Inc., Pleasanton, CA, USA) operated in electron-counted mode. Motion correction of images was done using the Alignframe function in IMOD^48^. Imaging was done using a Volta phase plate^49^ to increase contrast without high defocus, and the Gatan Imaging Filter (Gatan Inc., Pleasanton, CA, USA) with a slit width of 20 eV to increase contrast by removing inelastically scattered electrons^50^. After initially assessing cells at lower magnifications for suitability of ice thickness and plasma membrane integrity, tilt series were collected with a span of 100° (-50° to +50°; dose-symmetric scheme^51^) or 120° (-60° to +60°; dose-symmetric or bi-directional scheme) with 2° increments at a magnification of 33,000x (with a corresponding pixel size of 2.65 Å) and a defocus range of -1 to -4 μm. Each tilt series was collected with a cumulative dose of around 140 e^−^/ Å^2^. Once acquired, tilt series were aligned using the 10 nm colloidal gold as fiducials and reconstructed into tomograms by our in-house automated computation pipeline utilizing the IMOD software package^48^.

### Quantification, statistics, and reproducibility

We obtained a total of 218 *P. falciparum* tomograms (151 untreated and 67 GlyA-treated) from nine frozen grids prepared from six biological replicates (four untreated and two GlyA-treated) that were imaged over six multiple-day sessions. Each of the following quantifications is from a subset of 39 tomograms that resolved the feature of interest. We note that ∼50% of parasites imaged had a broken plasma membrane, resulting in a greater resolution of intracellular features compared with completely intact merozoites. We also note that parasites flattened on the grid, likely due to blotting. However, this flattening did not reflect on the shape of individual rhoptries, the AV, or the RSA. Flattening could have added subtle variations to the relative positions of these features, but their organizational patterns were evident despite the presence of such a potential effect.

#### Rhoptry distance from the AV

Shortest distance between the AV membrane and the rhoptry tip membrane for each rhoptry.

#### θ (orientation parameter for rhoptries at the AV)

Orientation of rhoptry 1 (θ_Rh1_) and rhoptry 2 (θ_Rh2_) necks with respect to the major axis of the AV (see below under ‘vesicle dimensions’ for more details). Angle was measured between the major axis of the AV and the line connecting the rhoptry tip membrane to the center point of the AV major axis.

#### Vesicle dimensions

The AV was approximated to an ellipsoid using only two axes (major and minor) instead of three for simplicity. The longest axis of each AV in 3-D was marked as the major axis (labeled as AV_maj_) while the shortest axis orthogonal to the major axis and intersecting it at the centroid was marked as the minor axis (labeled as AV_min_).

#### Eccentricity

(1-b^2^/a^2^)^1/2^, where ‘*a*’ is the semi-major axis, and ‘*b*’ is the semi-minor axis.

#### AV_dist_ (anchoring distance of AV)

Shortest distance between the AV membrane and the apex of the parasite plasma membrane, represented by the CD of the RSA.

#### Ψ (orientation parameter for AV)

Angle formed between the major axis of the AV and the line formed between the apex of the parasite plasma membrane and the centroid of the AV.

Quantifications were performed on IMOD-generated models. All model files were exported for analysis and plotting using Numpy, Matplotlib, and Seaborn libraries in Python 3.7. The corresponding graphs report the mean ± standard deviation values. For skewed distributions (θ and Ψ) we instead report the median value alone or boxplots displaying data quartiles.

### Pixel value analysis

To analyze the pixel values across line profiles through AVs, rhoptries, and RSAs, 3-D volumes were opened in IMOD and, for the AVs and rhoptries, 5 layers of voxels spanning a thickness of 5.3 nm were averaged along the Z axis. TIFF images of desired slices were subsequently imported into FIJI^52^ where a Gaussian filter was applied to smoothen the images. A line with a thickness of 10 pixels was drawn across the AVs and rhoptries, extending a few pixels beyond each membrane. Pixel values along these line profiles were exported for further analysis. For analysis of the RSA central channel, no filter was applied to the images in FIJI. Pixel values were inverted by subtracting each value from 255 so that electron dense regions appeared as peaks when plotted. Analysis and plotting of pixel values across each line were performed using Numpy, Matplotlib, and Seaborn libraries in Python 3.7.

### Rhoptry volume analysis

29 tomograms (10 parasites with an AV, 9 lacking the AV and with two distinct rhoptries, and 10 lacking the AV and with fused rhoptries) were analyzed in EMAN2^21, 22^ and UCSF Chimera^23^. Tomograms were imported into the EMAN2 tomogram annotation workflow and a neural network was trained to recognize and segment density corresponding to rhoptries within the tomogram. Segmented rhoptries were subsequently imported into UCSF Chimera, where the segmentations were cleaned up using volume eraser to erase false positive noise, a dome cap was maintained at the top and bottom of rhoptry segmentations following the rhoptry curvature to account for the missing wedge effect, and the volumes of the rhoptry segmentations were measured. Plotting of the rhoptry volumes was performed using Matplotlib and Seaborn libraries in Python 3.7.

### Generation of RSA subtomogram averages

Subtomogram alignment and averaging were performed using Dynamo^53, 54^. In IMOD, we performed manual inspections of merozoite tomograms at the apical end and identified RSA ultrastructures. Subtomogram boxes were centered in the middle of the RSA central channel and manually rotated to the same orientation among RSA ultrastructures. Model points and manual orientations were imported in Dynamo using custom scripts. Subtomogram averaging was initially performed using 149 RSAs without rotational symmetry. Densities with 8-fold rotational symmetry were seen throughout the RSA in the average, from the extracellular region to the AV (Supplementary Fig. 3). We therefore utilized this symmetry during particle picking, thus yielding 8 times more subtomograms, increasing the total number to 1,192 to further enhance the signal-to-noise ratio. The result was a better-resolved average that revealed a more elaborate ensemble of components than what was directly observed in the tomograms or from the unsymmetrized average.

RSAs from cells with different rhoptry morphologies (RSA-A: two rhoptries with an AV; RSA-2R: two rhoptries without an AV; RSA-1R: fused rhoptries without an AV) were manually sorted and then aligned and averaged independently from each other using Dynamo software (1,192 subtomograms from 149 RSA-As, 264 subtomograms from 33 RSA-2Rs, and 288 subtomograms from 36 RSA-1Rs). To generate an initial reference, each group of subtomograms was averaged using the manual orientations and without any computational alignment. An ellipsoid mask that covered the RSA and part of the plasma membrane was used in conjunction with each reference that was used to align all subtomograms. The alignment procedure started with a cone range (Euler angles Z and X) of 6° and an azimuth range (Euler angle Z’) of 22.5°, followed by 6 refinement iterations, each of which used half the angular search range of the previous iteration. A new reference was generated with the best two-thirds of aligned subtomograms, and the process was repeated two more times. This process was then repeated three more times with an initial azimuth range of 6° (in total, 6 iterations of alignment). We then incorporated 8-fold rotational symmetry of the RSAs by picking 8 subtomograms from each RSA particle (by iteratively rotating each particle by 45°) and performing another subtomogram alignment with cone and azimuth ranges of 6° for 3 iterations, yielding the final average for each kind of RSA (RSA-A, RSA-2R, and RSA-1R). Fourier shell correlation plots were calculated using the “Adaptive bandpass filtering” function in Dynamo.

### Figures presentation, modeling, and segmentations

Tomograms were oriented in 3-D using IMOD’s Slicer window such that the desired tomogram section was in view for presentation. To enhance contrast, 5-10 layers of voxels spanning thicknesses of 5.3-10.6 nm were averaged around the section of interest. Figures were prepared using Inkscape 1.1. Manual rhoptry segmentations were generated in IMOD and imported into ChimeraX^55^ for presentation. Segmentations of the RSA-A subtomogram average was done utilizing the Segger function in ChimeraX. Subsequent visualization and animations of the RSA were also generated using ChimeraX.

## Acknowledgements

We thank Dr. Stefan Steimle for his technical assistance with the Titan Krios G3i cryogenic electron microscope and the Singh Center for Nanotechnology and the Beckman Center for Cryo-Electron Microscopy at the University of Pennsylvania for hosting and supporting the use of the Titan Krios; Dana Hodge in the lab of A.O.J. for training in *P. falciparum* culture techniques; Liam Theveny from the lab of Y.-W.C. for providing subtomogram averages of the *T. gondii* and *C. parvum* RSAs; and other members of the labs of Y.-W.C., A.O.J., and M.L. for overall support and useful discussions. This work was supported in part by a David and Lucile Packard Fellowship for Science and Engineering (2019-69645) and a Pennsylvania Department of Health FY19 Health Research Formula Fund to Y.-W.C.; by the Mary L. and Matthew S. Santirocco College Alumni Society Undergraduate Research Grant to W.D.C.; by an EMBO fellowship (ALTF 58-2018) to A.G.; by an NIH/NIAID R01 AI103280, an R21 AI123808, an R21 AI130584, and an R61 DH105594 to A.O.J. who is an Investigator in the Pathogenesis of Infectious Diseases (PATH) of the Burroughs Wellcome Fund; and by a European Research Council advanced grant 833309 (KissAndSpitRhoptry) to M.L.

## Author contributions

M.M., M.L., and Y.-W.C. conceptualized and designed the experiments. M.M. cultured and isolated parasites provided by A.O.J. M.M.C. provided a protocol and consultations for the efficient isolation of merozoites. P.M. provided further training and useful insights towards *P. falciparum* culture and merozoite isolation. M.M. prepared frozen grids and performed cryo-ET, with training from S.K.M., using an automated data-processing pipeline for on-the-fly tomogram reconstruction that was established by W.D.C., who also provided additional computational support during data collection, processing, and management. M.M. analyzed the tomograms, performed subtomogram averaging, and analyzed the RSA structure. W.D.C. performed data analysis of rhoptry volumes. S.K.M., A.G., M.M.C., and M.L. provided important insights for the interpretation of data. M.M. prepared the manuscript with critical inputs and revisions from all authors.

## Competing interests

The authors have no competing financial interests.

**Supplementary Information** is available for this paper.

---

**Supplementary Figure 1.**
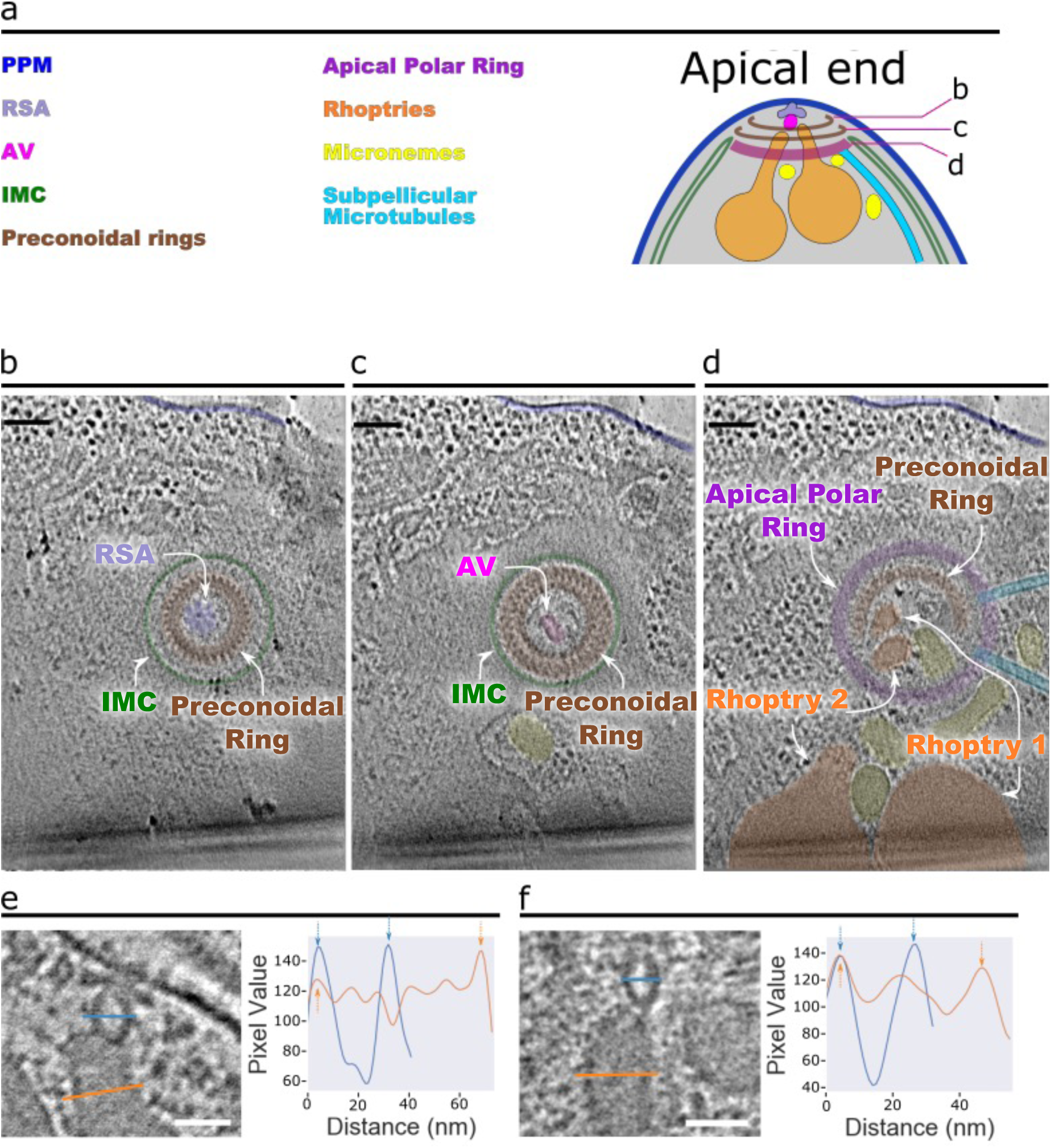
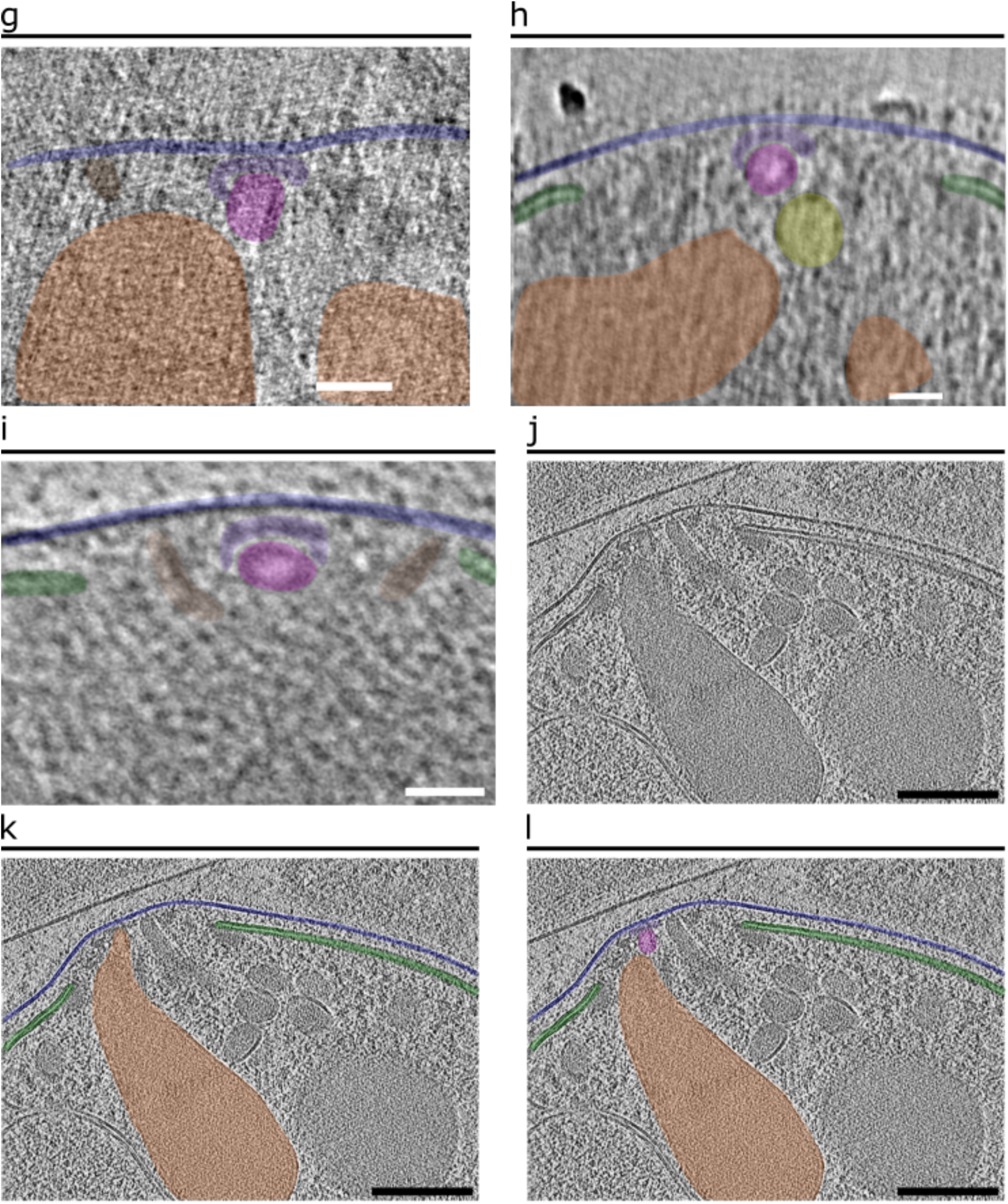
The apical complex and apical vesicle of *P. falciparum* merozoites. **(a)** Simplified schematic of a *P. falciparum* merozoite apical region. **(b–d)** 2-D slices through a tomogram of a top-down view of the merozoite apical complex with color overlays of all the observed apical end components: apical vesicle (AV; pink), parasite plasma membrane (PPM; blue), rhoptry (orange), rhoptry secretory apparatus (RSA; light blue), microneme (yellow), inner membrane complex (IMC; green), preconoidal rings (brown), apical polar ring (purple), and subpellicular microtubules (cyan). The cross-sectional planes used in b-d are arranged such that they start at the parasite apex and move inwards into the cell. Scale bar = 100 nm. **(e, f)** Additional examples of 2-D slices of the AV and rhoptry (left). Blue and orange lines across the AV and rhoptry, respectively, denote the locations of pixel values plotted on the right. Arrows in the pixel plots point to peaks in values corresponding to membranes. Scale bars = 50 nm. **(g–i)** 2-D slices from tomograms showing either one rhoptry (g, h) or no rhoptries (i) docked at the AV. Scale bars = 50 nm. **(j)** 2-D slice from a *P. falciparum* merozoite apical end tomogram within a mature schizont, adapted from [30]. **(k)** The same 2-D slice from panel j with the original color overlay of the rhoptry of interest, according to [30]. **(l)** The same 2-D slice from panel j with an updated color overlay of the rhoptry of interest, the AV, and the RSA, based on our analysis. Scale bars = 200 nm.

**Supplementary Figure 2.**
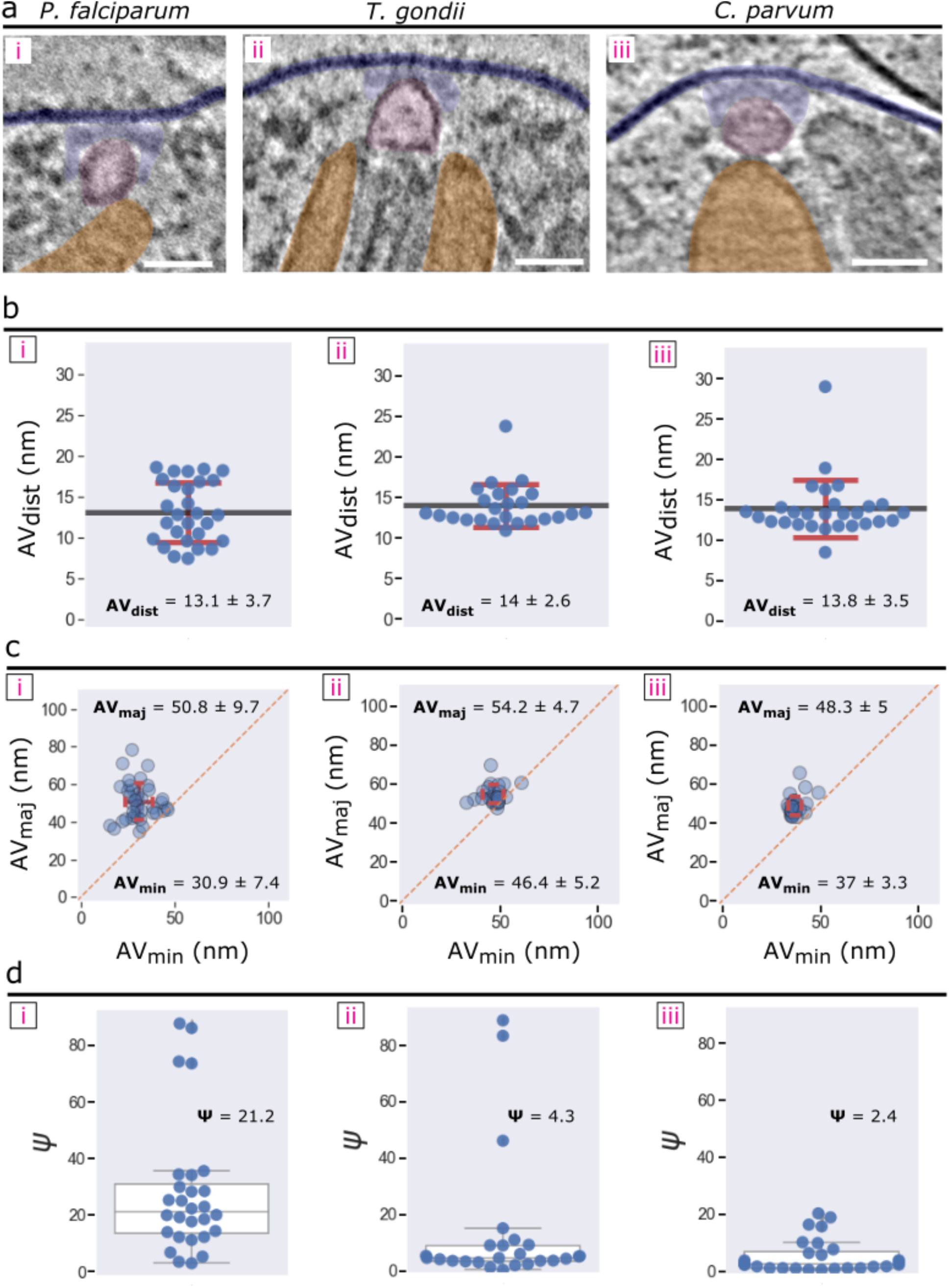
Comparison of the apical vesicles from *P. falciparum*, *T. gondii*, and *C. parvum*. **(a)** Representative 2-D slices of tomograms displaying the AV from each organism (left – *P. falciparum*; middle – *T. gondii*; and right – *C. parvum*), highlighting the PPM (dark blue), RSA (light blue), AV (pink), and rhoptries (orange). Scale bars = 50 nm. **(b)** Plots of the shortest distance (AV_dist_) between the PPM apex and the AV membrane from several cells each organism, showing the mean ± sd. **(c)** Plots of the major (AV_maj_) versus minor axis (AV_min_) of the AV from each organism, showing mean ± sd. **(d)** Plots of the angle (Ψ) at which the major axis of the AV is oriented with respect to the line connecting the PPM apex and the AV centroid each organism. The median value is shown. N = 39 AVs for *P. falciparum*, 25 AVs for *T. gondii*, and 28 AVs for *C. parvum*.

**Supplementary Figure 3.**
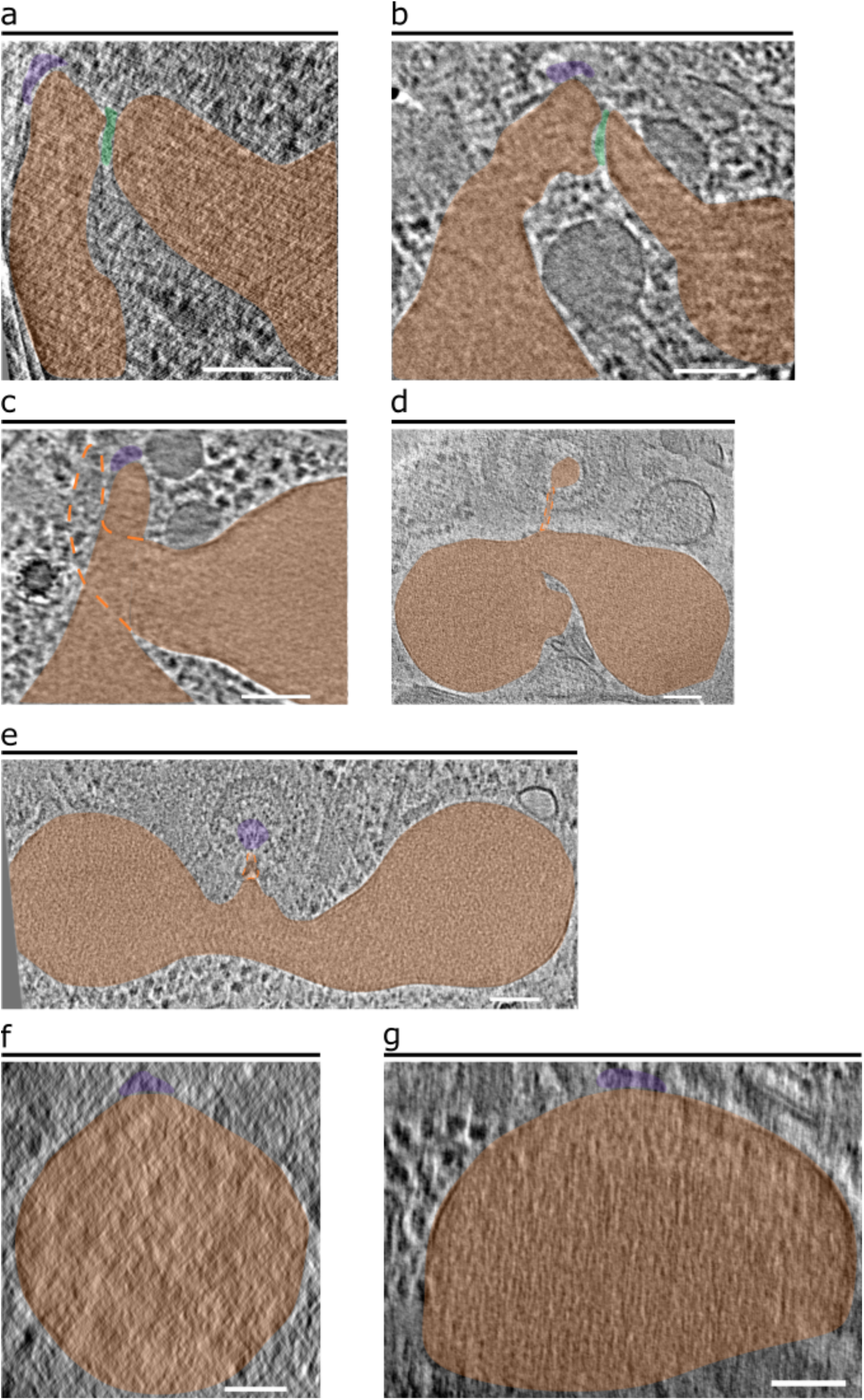
Various fusion states of the *P. falciparum* rhoptries. **(a–c)** 2-D slices through tomograms of apical ends showing the absence of the apical vesicle at the rhoptry tips and either bulging of one rhoptry towards the other (a, b) or twisting of the rhoptries (c). Dashed orange lines in panel c denote the portion of the rhoptry neck that is out of view. A mesh of electron-dense material (green) can be observed between rhoptries (orange) that display bulging towards the other. One rhoptry is docked directly at the RSA (purple). **(d, e)** Examples in which the two rhoptries are fused together and docked directly at the RSA. **(f, g)** Examples in which only one, very large rhoptry with no discernable neck region was observed and docked directly at the RSA. Scale bars = 100 nm.

**Supplementary Figure 4.**
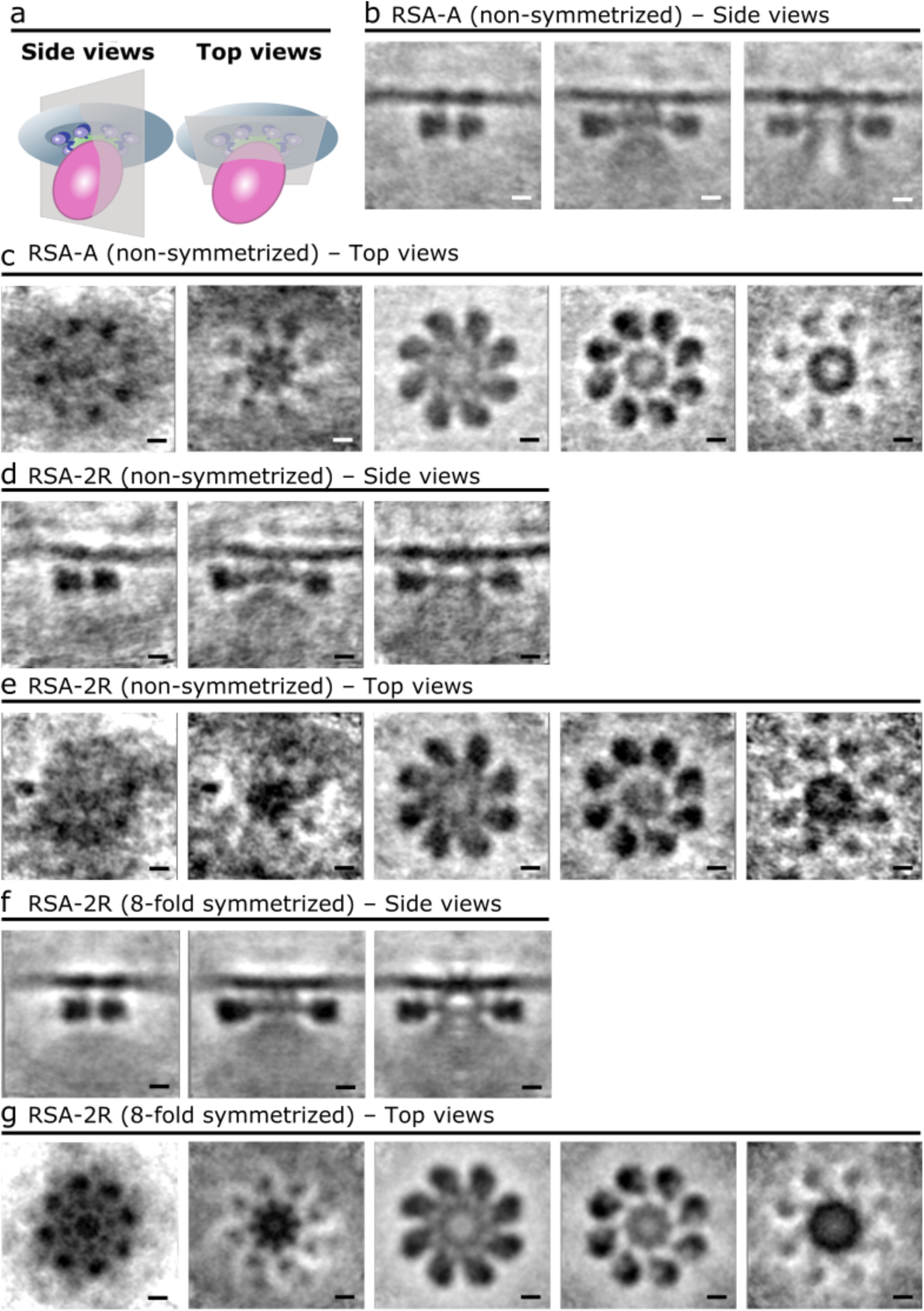
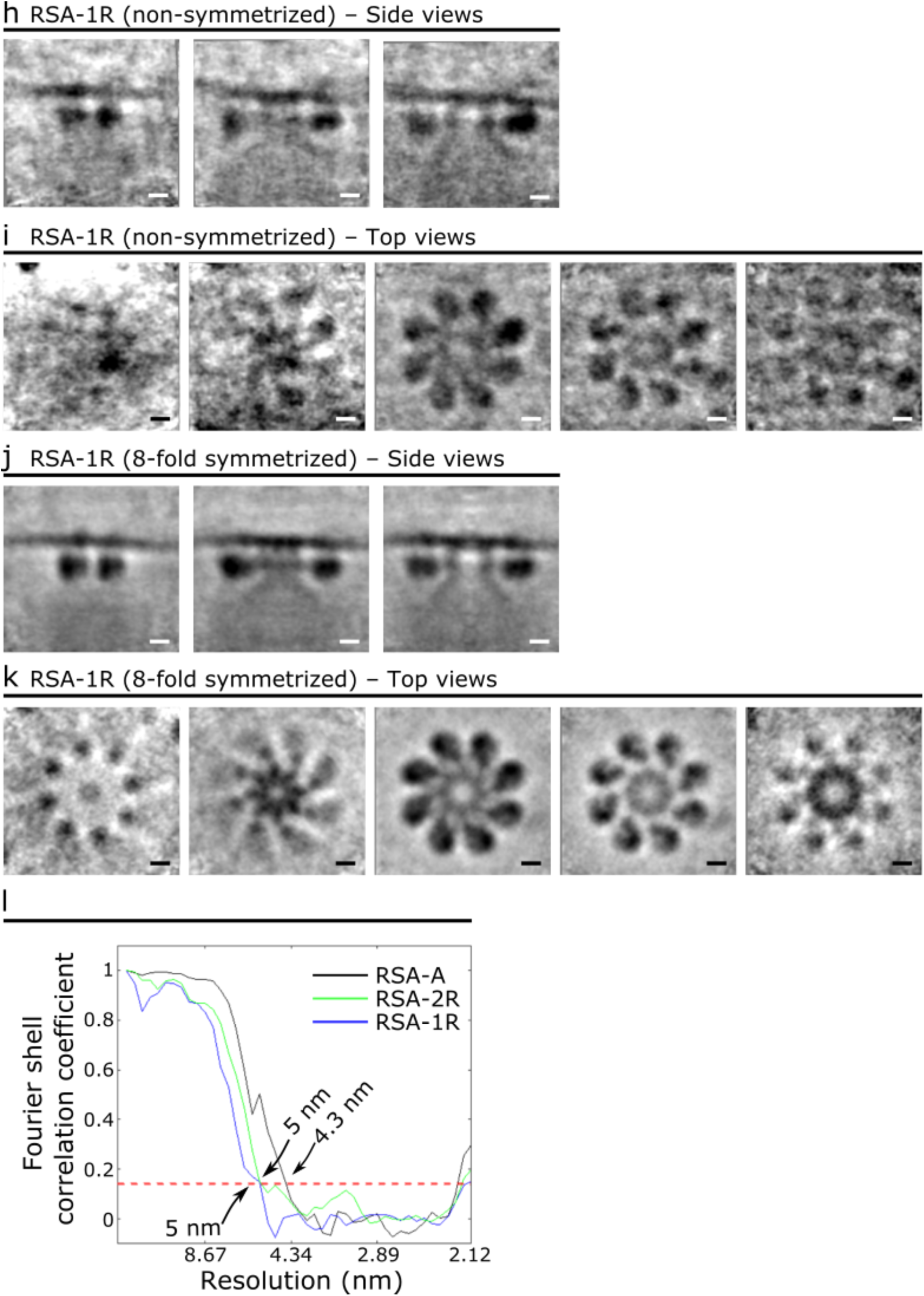
Subtomogram averages of the *P. falciparum* RSA. **(a)** Schematic of the RSA, apical vesicle, and PPM demonstrating the top view and side view orientations used to generate 2-D slices of the subtomogram averages. **(b, c)** Side views (b) and top view (c) of the subtomogram average of the RSA-A structure before applying 8-fold symmetry. Left to right panels: from peripheral to central cross sections (b) or from extracellular to intracellular cross sections (c). **(d**–**g)** Side views (d, f) and top view (e, g) of the subtomogram average of the RSA-2R structure, before and after applying 8-fold symmetry. **(h**–**k)** Side views (h, j) and top views (i, k) of the subtomogram average of the RSA-1R structure, before and after applying 8-fold symmetry. **(i)** Gold standard Fourier shell correlation plot of the final, 8-fold symmetrized subtomogram averages of the three RSA structures. Scale bars = 10 nm.

**Supplementary Figure 5.**
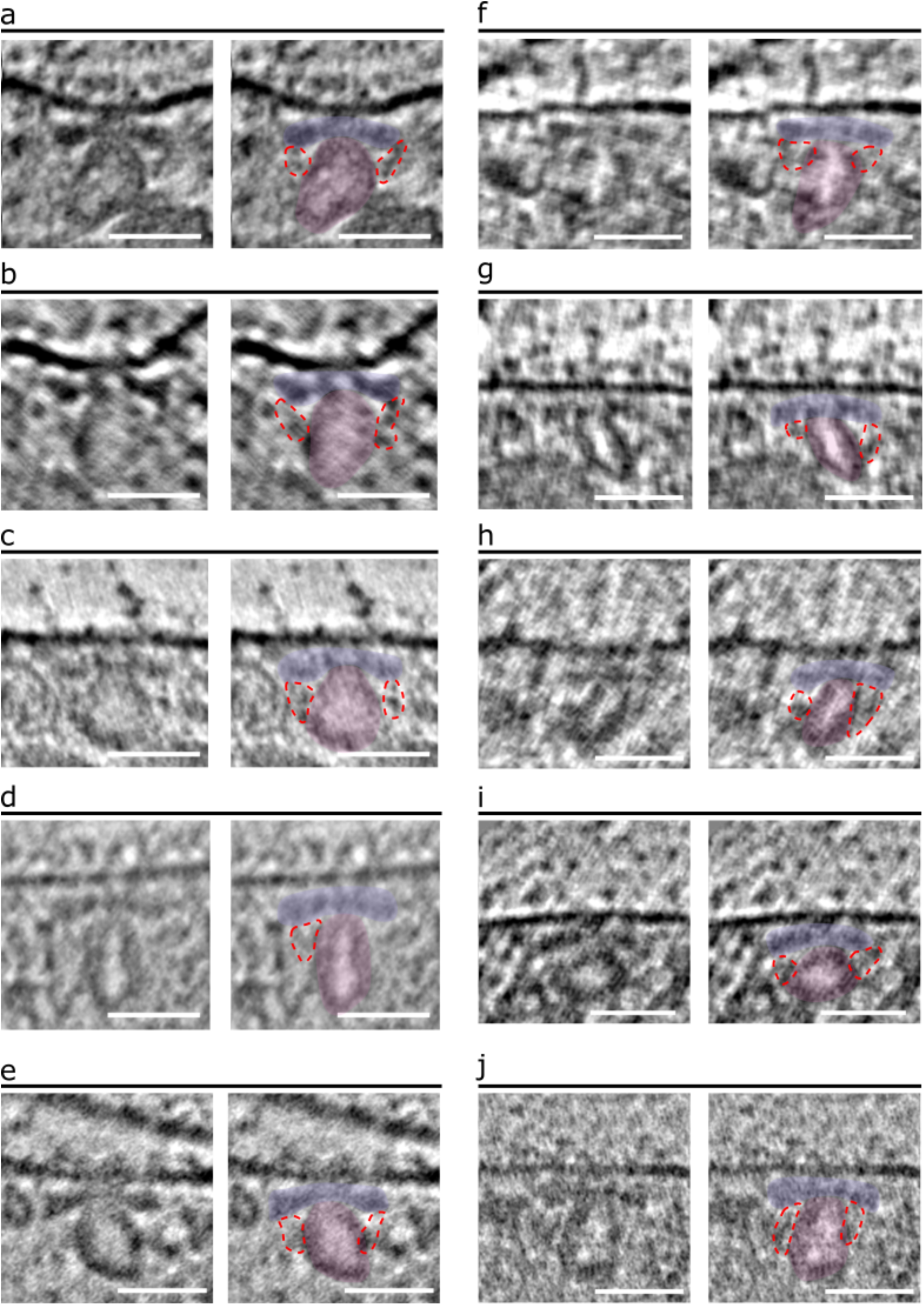
Representative cryo-ET images of the *P. falciparum* AV and RSA. **(a-j)** 2-D slices with of merozoite apexes (left panels) with color overlays (right panels) highlighting the RSA (light blue), the AV (pink), and the connections from the RSA A-II/III densities to the AV (encircled in red dashed lines). Scale bars = 50 nm.

**Movie 1.** Subtomogram average of the *P. falciparum* RSA-A structure slicing first through the top view from extracellular to intracellular and back, followed by slicing through the side views.

**Movie 2.** 3-Dimensional view of the subtomogram average of the *P. falciparum* RSA-A structure.

**Movie 3.** Subtomogram average of the *P. falciparum* RSA-2R structure slicing first through the top view from extracellular to intracellular and back, followed by slicing through the side views.

**Movie 4.** Subtomogram average of the *P. falciparum* RSA-1R structure slicing first through the top view from extracellular to intracellular and back, followed by slicing through the side views.

**Movie 5.** 3-Dimensional view of the *P. falciparum* RSA structure, highlighting conformational changes between the RSA-A and RSA-1R states.

## References

1. World malaria report 2021. (2021).

2. World Malaria Report 2020: 20 years of global progress and challenges. (2020).

3. Cowman, A. F., Healer, J., Marapana, D. & Marsh, K. Malaria: Biology and Disease. Cell 167, 610–624 (2016).

4. Blader, I. J., Coleman, B. I., Chen, C.-T. & Gubbels, M.-J. Lytic Cycle of Toxoplasma gondii: 15 Years Later. Annu. Rev. Microbiol 69, 463–485 (2015).

5. Guérin, A. & Striepen, B. The Biology of the Intestinal Intracellular Parasite Cryptosporidium. Cell Host Microbe 28, 509–515 (2020).

6. Dubois, D. J. & Soldati-Favre, D. Biogenesis and secretion of micronemes in Toxoplasma gondii. Cell. Microbiol. 21, e13018 (2019).

7. Ben Chaabene, R., Lentini, G. & Soldati-Favre, D. Biogenesis and discharge of the rhoptries: key organelles for entry and hijack of host cells by the Apicomplexa. Mol. Microbiol. (2020). doi:10.1111/mmi.14674

8. Cowman, A. F., Tonkin, C. J., Tham, W.-H. & Duraisingh, M. T. The Molecular Basis of Erythrocyte Invasion by Malaria Parasites. Cell Host Microbe 232–245 (2017). doi:10.1016/j.chom.2017.07.003

9. Riglar, D. T. et al. Super-resolution dissection of coordinated events during malaria parasite invasion of the human erythrocyte. Cell Host Microbe 9, 9–20 (2011).

10. Sparvoli, D. & Lebrun, M. Unraveling the Elusive Rhoptry Exocytic Mechanism of Apicomplexa. Trends Parasitol. (2021). doi:10.1016/j.pt.2021.04.011

11. Hanssen, E. et al. Electron tomography of Plasmodium falciparum merozoites reveals core cellular events that underpin erythrocyte invasion. Cell. Microbiol. 15, 1457–1472 (2013).

12. Aikawa, M., Miller, L. H., Johnson, J. & Rabbege, J. Erythrocyte entry by malarial parasites. A moving junction between erythrocyte and parasite. J. Cell Biol. 77, 72–82 (1978).

13. Miller, L. H., Aikawa, M., Johnson, J. G. & Shiroishi, T. Interaction between cytochalasin B-treated malarial parasites and erythrocytes. J. Exp. Med. 149, 172–184 (1979).

14. Bannister. L. H., Mitchell, G. H., Butcher, G. A. & Dennis, E. D. Lamellar Membranes Associated With Rhoptries In Erythrocytic Merozoites Of Plasmodium Knowlesi: A Clue To The Mechanism Of Invasion. Parasitology 92, 291–303 (1986).

15. Burrell, A. et al. Cellular electron tomography of the apical complex in the apicomplexan parasite Eimeria tenella shows a highly organised gateway for regulated secretion. bioRxiv (2021). doi:10.1101/2021.06.17.448283

16. Singh, S., Alam, M. M., Pal-Bhowmick, I., Brzostowski, J. A. & Chitnis, C. E. Distinct External Signals Trigger Sequential Release of Apical Organelles during Erythrocyte Invasion by Malaria Parasites. PLoS Pathog 6, e1000746 (2010).

17. Aquilini, E. et al. An Alveolata secretory machinery adapted to parasite host cell invasion. Nat. Microbiol. (2021). doi:10.1038/s41564-020-00854-z

18. Weiss, G. E. et al. Revealing the Sequence and Resulting Cellular Morphology of Receptor-Ligand Interactions during Plasmodium falciparum Invasion of Erythrocytes. PLoS Pathog. 11, e1004670 (2015).

19. Volz, J. C. et al. Essential Role of the PfRh5/PfRipr/CyRPA Complex during Plasmodium falciparum Invasion of Erythrocytes. Cell Host Microbe 20, 60–71 (2016).

20. Mageswaran, S. K. et al. In situ ultrastructures of two evolutionarily distant apicomplexan rhoptry secretion systems. Nat. Commun. 12, 1–12 (2021).

21. Tang, G. et al. EMAN2: An extensible image processing suite for electron microscopy. J. Struct. Biol. 157, 38–46 (2007).

22. Chen, M. et al. Convolutional neural networks for automated annotation of cellular cryo-electron tomograms. Nat. Methods 14, 983–985 (2017).

23. Pettersen, E. F. et al. UCSF Chimera—A visualization system for exploratory research and analysis. J. Comput. Chem. 25, 1605–1612 (2004).

24. Dubremetz, J.-F. & Entzeroth, R. Exocytic events during cell invasion by Apicomplexa. in Advances in Cellular and Molecular Biology of Membranes and Organelles (ed. Plattner, H.) 83–98 (1994).

25. Dubremetz, J. F. & Torpier, G. Freeze fracture study of the pellicle of an Eimerian sporozoite (Protozoa, Coccidia). J. Ultrastruct. Res. 62, 94–109 (1978).

26. Porchet, E. & Torpier, G. [Freeze fracture study of Toxoplasma and Sarcocystis infective stages (author’s transl)]. Z Parasitenkd 54, 101–124 (1977).

27. Kudryashev, M. et al. Structural basis for chirality and directional motility of Plasmodium sporozoites. Cell. Microbiol. 14, 1757–1768 (2012).

28. Anton, L., Cobb, D. W. & Ho, C.-M. Structural parasitology of the malaria parasite Plasmodium falciparum. Trends Biochem. Sci. (2021). doi:10.1016/J.TIBS.2021.10.006

29. Sun, S. Y. et al. Cryo-ET reveals two major tubulin-based cytoskeleton structures in Toxoplasma gondii. bioRxiv (2021). doi:10.1101/2021.05.23.445366

30. Bisson, C., Hecksel, C. W., Gilchrist, J. B. & Fleck, R. A. Preparing Lamellae from Vitreous Biological Samples using a Dual-Beam Scanning Electron Microscope for Cryo-Electron Tomography . J. Vis. Exp e62350 (2021). doi:10.3791/62350

31. Counihan, N. A., Kalanon, M., Coppel, R. L. & De Koning-Ward, T. F. Plasmodium rhoptry proteins: why order is important. Trends Parasitol. 29, (2013).

32. Liffner, B. et al. PfCERLI1 is a conserved rhoptry associated protein essential for Plasmodium falciparum merozoite invasion of erythrocytes. Nat. Commun. 11, (2020).

33. Tetley, L., Brown, S. M. A., Mcdonald, V. & Coombsl, G. H. Ultrastructural analysis of the sporozoite of Cryptosporidium parvum. Microbiology 144, 3249–3255 (1998).

34. Paredes-Santos, T. C., De Souza, W. & Attias, M. Dynamics and 3D organization of secretory organelles of Toxoplasma gondii. J. Struct. Biol. 177, 420–430 (2012).

35. Dubremetz, J. F. Rhoptries are major players in Toxoplasma gondii invasion and host cell interaction. Cell. Microbiol. 9, 841–848 (2007).

36. Plattner, H. Trichocysts-Paramecium’s Projectile-like Secretory Organelles: Reappraisal of their Biogenesis, Composition, Intracellular Transport, and Possible Functions. J. Eukaryot. Microbiol. 64, 106–133 (2017).

37. Lyth, O. et al. Cellular dissection of malaria parasite invasion of human erythrocytes using viable Plasmodium knowlesi merozoites. Sci. Rep. 8, 10165 (2018).

38. Aikawa, M., Miller, L. H., Rabbege, J. R. & Epstein, N. Freeze-Fracture Study on the Erythrocyte Membrane during Malarial Parasite Invasion. J. Cell Biol. 91, 55–62 (1981).

39. Liffner, B., Balbin, J. M., Wichers, J. S., Gilberger, T.-W. & Wilson, D. W. The Ins and Outs of Plasmodium Rhoptries, Focusing on the Cytosolic Side. Trends Parasitol. (2021). doi:10.1016/J.PT.2021.03.006

40. Nichols, B. A., Chiappino, M. L. & Richard O’connor, G. Secretion from the Rhoptries of Toxoplasma gondii during Host-Cell Invasion. J. Ultrastruct. Res. 83, 85–98 (1983).

41. Baum, J. et al. Reticulocyte-binding protein homologue 5 – An essential adhesin involved in invasion of human erythrocytes by Plasmodium falciparum. Int. J. Parasitol. 39, 371– 380 (2009).

42. de Oliveira, L. S. et al. Calcium in the Backstage of Malaria Parasite Biology. Front. Cell. Infect. Microbiol. 11, (2021).

43. Suarez, C. et al. A lipid-binding protein mediates rhoptry discharge and invasion in Plasmodium falciparum and Toxoplasma gondii parasites. Nat. Commun. 10, (2019).

44. Boyle, M. J. et al. Isolation of viable Plasmodium falciparum merozoites to define erythrocyte invasion events and advance vaccine and drug development. PNAS 107, 14378–14383 (2010).

45. Iancu, C. V. et al. Electron cryotomography sample preparation using the Vitrobot. Nat. Protoc. 1, 2813–2819 (2006).

46. Mastronarde, D. N. Automated electron microscope tomography using robust prediction of specimen movements. J. Struct. Biol. 152, 36–51 (2005).

47. Xuong, N. H. et al. Future Directions for Camera Systems in Electron Microscopy. Methods Cell Biol. 79, 721–739 (2007).

48. Kremer, J. R., Mastronarde, D. N. & McIntosh, J. R. Computer visualization of three-dimensional image data using IMOD. J. Struct. Biol. 116, 71–76 (1996).

49. Danev, R., Buijsse, B., Khoshouei, M., Plitzko, J. M. & Baumeister, W. Volta potential phase plate for in-focus phase contrast transmission electron microscopy. Proc. Natl. Acad. Sci. U. S. A. 111, 15635–15640 (2014).

50. Krivanek, O. L., Friedman, S. L., Gubbens, A. J. & Kraus, B. An imaging filter for biological applications. Ultramicroscopy 59, 267–282 (1995).

51. Turoňová, B. et al. Benchmarking tomographic acquisition schemes for high-resolution structural biology. Nat. Commun. 11, (2020).

52. Schindelin, J. et al. Fiji: an open-source platform for biological-image analysis. Nat. Methods 9, 676–682 (2012).

53. Castaño-Díez, D., Kudryashev, M., Arheit, M. & Stahlberg, H. Dynamo: A flexible, user-friendly development tool for subtomogram averaging of cryo-EM data in high-performance computing environments. J. Struct. Biol. 178, 139–151 (2012).

54. Castaño-Díez, D. The Dynamo package for tomography and subtomogram averaging: components for MATLAB, GPU computing and EC2 Amazon Web Services. Acta Cryst. D73, 478–487 (2017).

55. Pettersen, E. F. et al. UCSF ChimeraX: Structure visualization for researchers, educators, and developers. Protein Sci. 30, 70–82 (2021).

